# Single-cell multiome uncovers differences in glycogen metabolism underlying species-specific speed of development

**DOI:** 10.1101/2024.09.03.610938

**Authors:** Alexandra de la Porte, Julia Schröder, Moritz Thomas, Johanna Geuder, Michael Sterr, Xavier Pastor, Leslie E. Sanderson, Tahsin Stefan Barakat, Wolfgang Enard, Carsten Marr, Christian Schröter, Micha Drukker

## Abstract

Embryos from different mammalian species develop at characteristic timescales. These timescales are recapitulated during the differentiation of pluripotent stem cells *in vitro*. Specific genes and molecular pathways that modulate cell differentiation speed between mammalian species remain to be determined. Here we use single-cell multi-omic analysis of neural differentiation of mouse, cynomolgus and human pluripotent cells to identify regulators for differentiation speed. We demonstrate that species-specific transcriptome dynamics are mirrored at the chromatin level, but that the speed of neural differentiation is insensitive to manipulations of cell growth and cycling. Exploiting the single-cell resolution of our data, we identify glycogen storage levels regulated by UDP-glucose pyrophosphorylase 2 (UGP2) as a species-dependent trait of pluripotent cells, and show that lowered glycogen storage in UGP2 mutant cells is associated with accelerated neural differentiation. The control of energy storage could be a general strategy for the regulation of cell differentiation speed.

## Introduction

Mammalian embryonic development adheres to a strict sequence of events, yet the speed of development varies significantly among species: while it takes 13 days for human embryos to progress from oocyte fertilization to gastrulation, it only takes six days for mice to reach this stage^1^. Similarly, the timescales for organogenesis and later of neuronal differentiation, both in the peripheral nervous system and midbrain, are considerably longer in humans compared to mice^2^ (**Figure 1A**). Identifying the genetic and physiological basis for species-specific developmental timing has become an area of intense research in the past years, but specific genetic or gene-regulatory differences that underlie this phenomenon still remain to be reported.

**Figure 1:**
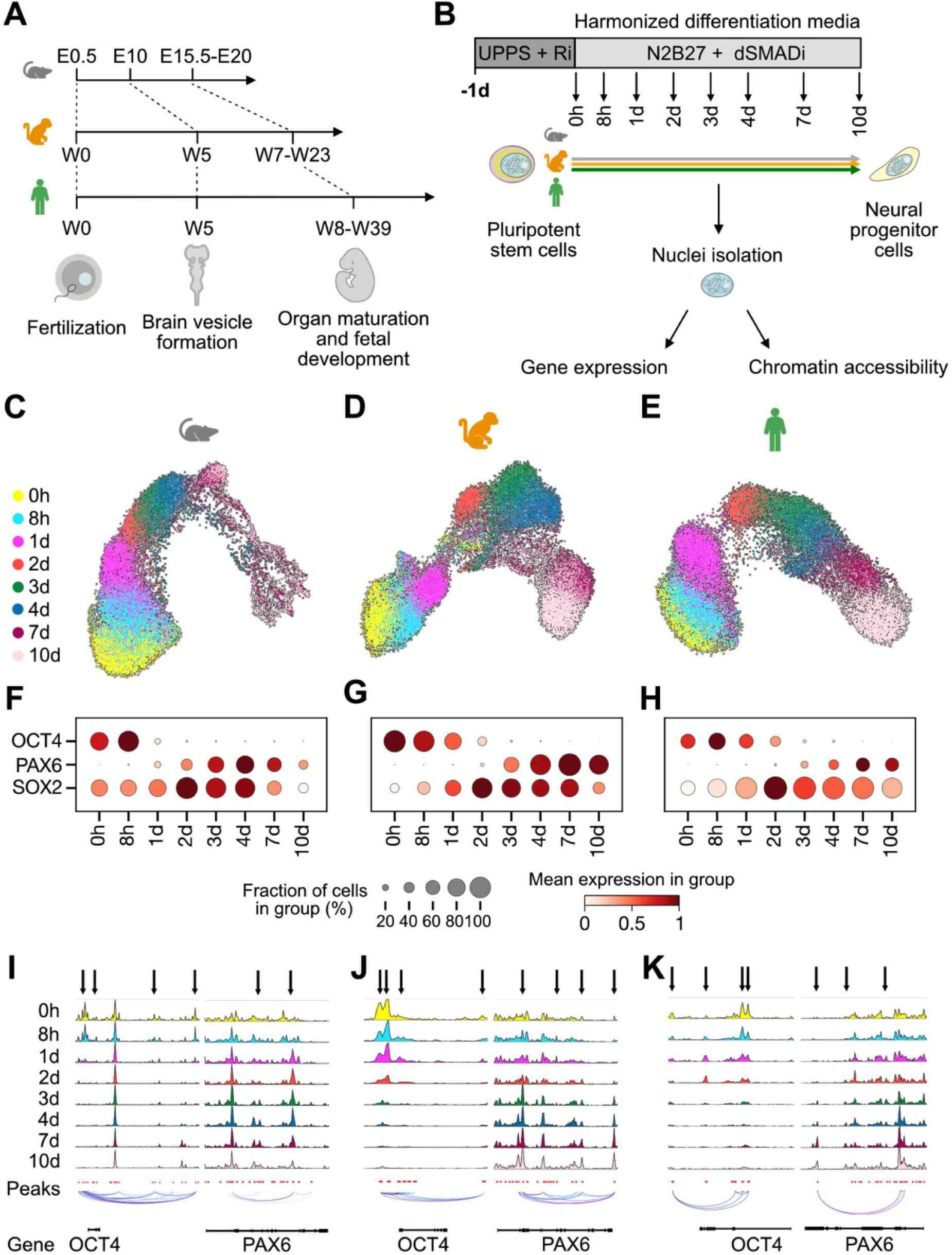
Single-cell multiome sequencing captures species-specific differences in neural differentiation. A. Embryonic developmental speed differences between mouse (measured in embryonic days, E), cynomolgus, and human (measured in weeks, W). B. Schematic of neural progenitor differentiation of pluripotent stem cells from the three species in a harmonized medium (N2B27 + dSMADi). Arrows indicate 8 time points chosen for single cell gene expression and chromatin accessibility measurements within the first 10 days. C-D. Two-dimensional UMAP embedding of mouse (C), cynomolgus (D) and human (E) RNA expression in 24,157, 23,914 and 26,043 single cells, respectively, during NPC differentiation. F-H. Expression of selected marker genes indicates faster transition from a stem cell state (represented by OCT4 and SOX2 expression) to a neural progenitor state (PAX6 and SOX2 expression), in mouse (F) compared to cynomolgus (G) and human (H). I-K. Changes in local chromatin accessibility of the stem cell marker OCT4 (left) and the neural progenitor state marker PAX6 (right) during 10 day differentiation in mouse (I), cynomolgus (J) and human (K) cells. Peaks of interest are indicated by arrows.

*In vivo* developmental pace is mirrored by the speed of *in vitro* differentiation of pluripotent stem cells (PSCs), indicating that cell differentiation speed has a cell-intrinsic genetic basis^3–6^. PSCs can either be derived directly from embryos, or generated as induced pluripotent stem cells (iPSCs) from the tissues of various mammalian species^7–11^. When cultured under appropriate conditions, these cells closely resemble the state of the pluripotent epiblast just before the beginning of gastrulation. This primed pluripotent state captured by human embryonic stem cells (hESCs), mouse epiblast stem cells (mEpiSCs), and iPSCs from both human and non-human primates^7,8,12^ can serve as a useful common starting point for comparative studies.

Previous studies using PSCs to investigate mechanisms of differentiation speed have to a large degree focused on comparisons between human and mouse cells, and only recently have a broader range of species been included^13^. A key challenge in such comparative studies is that PSCs are commonly established and differentiated in species-specific maintenance and differentiation media. While these media formulations may maximize the viability of cells from each species, they leave open the possibility that species-specific differentiation speed is at least in part caused by differences in external metabolic and signaling environments. External signaling through factors produced by the cells themselves is an additional candidate mechanism for regulating differentiation speed in lineages that are specified in response to constant paracrine signaling, such as the mesoderm or the cells of the peripheral nervous system.

Irrespective of these challenges, recent studies have started to implicate diverse general physiological cellular traits in the control of species-specific differentiation speed, such as differential protein stability^6,14^, biochemical reaction rates^5,13^, and mitochondrial activity^4,15^. Whether the same physiological parameters regulate differentiation speed of different lineages is however an open question. It is also not known at which level of regulation - chromatin, gene expression or post-transcriptional and post-translational events - species-specific differentiation speeds emerge. Finally, the evolutionary changes in cell physiology that underlie altered differentiation speed must have arisen from genetic changes that determine either the activity or the expression magnitude of specific proteins in the cell. In the context of differentiation speed, there are to date only few examples for specific species-specific changes to protein activity.

Cell differentiation *in vitro* is a heterogeneous process, in which divergent cellular dynamics lead to differences between individual cells. Recently, it has become possible to access this cellular diversity and dynamics at several levels of regulation with single-cell multiomic sequencing technologies^16^. Multiomic sequencing simultaneously profiles gene expression and chromatin accessibility in single cells, thereby enabling the reconstruction of developmental lineage trajectories which can ultimately inform temporal regulatory dependencies^17^.

In this study, we established identical stem cell culture and neuroectoderm differentiation modalities for hESCs, mEpiSCs and cynomolgus iPSC (cyiPSCs), and applied time-resolved multiomic single-cell profiling during neural progenitor differentiation to unravel the mechanisms underlying species-specific differentiation speed^18,19^. Through combined single-cell gene expression (scRNA-seq) and chromatin accessibility analysis (scATAC-seq), we show that developmental speed differences are governed by chromatin dynamics. Employing metabolic interventions, we showed that differentiation dynamics of neural progenitors can be uncoupled from anabolic processes. Finally, we developed strategies to identify candidate regulators of differentiation speed based on our time-resolved single-cell data. Through his approach, we identified glucose storage through UDP-glucose pyrophosphorylase 2 (UGP2) as a mechanism for the species-specific regulation of differentiation speed.

## Results

### Single-cell multiome sequencing for comparative analysis of species-specific differentiation speeds under harmonized culture and differentiation conditions

To investigate the speed of cell differentiation during anterior neural development (**Figure 1A**), we utilized epiblast-like primed-state hESCs, mEpiSCs, and cyiPSCs to perform comparative *in vitro* differentiation. To standardize culture conditions for the different species and thus eliminate external influences, the cells were gradually adapted to a unified medium and passaged at least twice before further experiments (**Figure S1A**). We identified Universal Primate Pluripotent Stem Cell Media (UPPS) reported by Stauske et al.^19^ as the ideal harmonized medium, confirmed by colony morphology (**Figure S1B**), and by the homogeneous expression of pluripotency markers in flow cytometry (**Figure S1C**) and immunofluorescence imaging (**Figure S1D**).

Next, we adapted a common protocol for neural progenitor cell (NPC) differentiation through dual SMAD inhibition (dSMADi, see Methods for details)^18^. In all three species, we observed quick downregulation of pluripotency-related markers OCT4 (also known as POU5F1) and NANOG, and upregulation of neural markers SOX1 and PAX6, both in immunofluorescence imaging (**Figure S1E**) and quantitative reverse transcription PCR (RT-qPCR) (**Figure S1F**). Overall, our comprehensive examination indicated successful pluripotent stem cell maintenance and NPC differentiation across all cell lines using standardized protocols.

To compare the differentiation speed of the three species in detail, we employed single-cell multiome sequencing, combining gene expression and chromatin accessibility sequencing. For each species, we conducted NPC differentiation over a ten-day time course, collecting samples at 0 hours, 8 hours, 1 day, 2 days, 3 days, 4 days, 7 days, and 10 days, to capture early changes in transcription and chromatin accessibility (**Figure 1B**). To minimize sequencing batch effects, we combined cells from each time point of all three species for nuclei isolation and subsequent scRNA-seq and scATAC-seq library preparation. For demultiplexing, single-cell multiome data was aligned against each genome separately, and each cell was assigned to the respective species (see Methods for details). UMAP plots derived from scRNA-seq of differentiating cells accurately depicted the linear path of NPC differentiation (**Figures 1C-E**). When examining the expression of marker genes, the observed dynamics aligned with the known differentiation trajectory (**Figures 1F-H**). For example, pluripotency marker OCT4 gradually decreased during NPC differentiation, while the neural marker PAX6 exhibited a gradual increase in expression with distinct, species-specific dynamics. The scATAC-seq data likewise provided evidence of a linear differentiation process, illustrated by the progressive reduction of accessible chromatin at the OCT4 gene locus and increase of accessible chromatin at the PAX6 gene locus, corresponding to the respective decrease and increase in gene expression (**Figure 1I-K**).

### Quantification of species-specific differentiation rates from single-cell multiome data

To examine the speed of differentiation across pluripotent stem cells among the three species, we created a unified cross-species representation by mapping cells from human and cynomolgus onto the mouse reference embedding using Scanpy’s ingest tool^20^ (**Figure S2A**). The combined UMAP visualization highlighted that cynomolgus cells were slightly further in the differentiation trajectory compared to human cells at each time point, indicating a marginally faster differentiation (**Figure S2B**). More specifically, cynomolgus cells sampled at day 10 mapped to mouse cells at day 3-7 (**Figure 2A**). A more pronounced deviation was seen with human cells sampled at day 10, which mapped to mouse cells from day 3 and day 4 (**Figure 2B**). We quantified the differentiation speed relative to mouse cells using a linear regression model, fitted through each mapped time point for human and cynomolgus cells, and found that that mouse cells differentiated on average 2.2 times faster than cynomolgus cells, and 2.4 times faster than human cells (**Figure 2C**).

**Figure 2:**
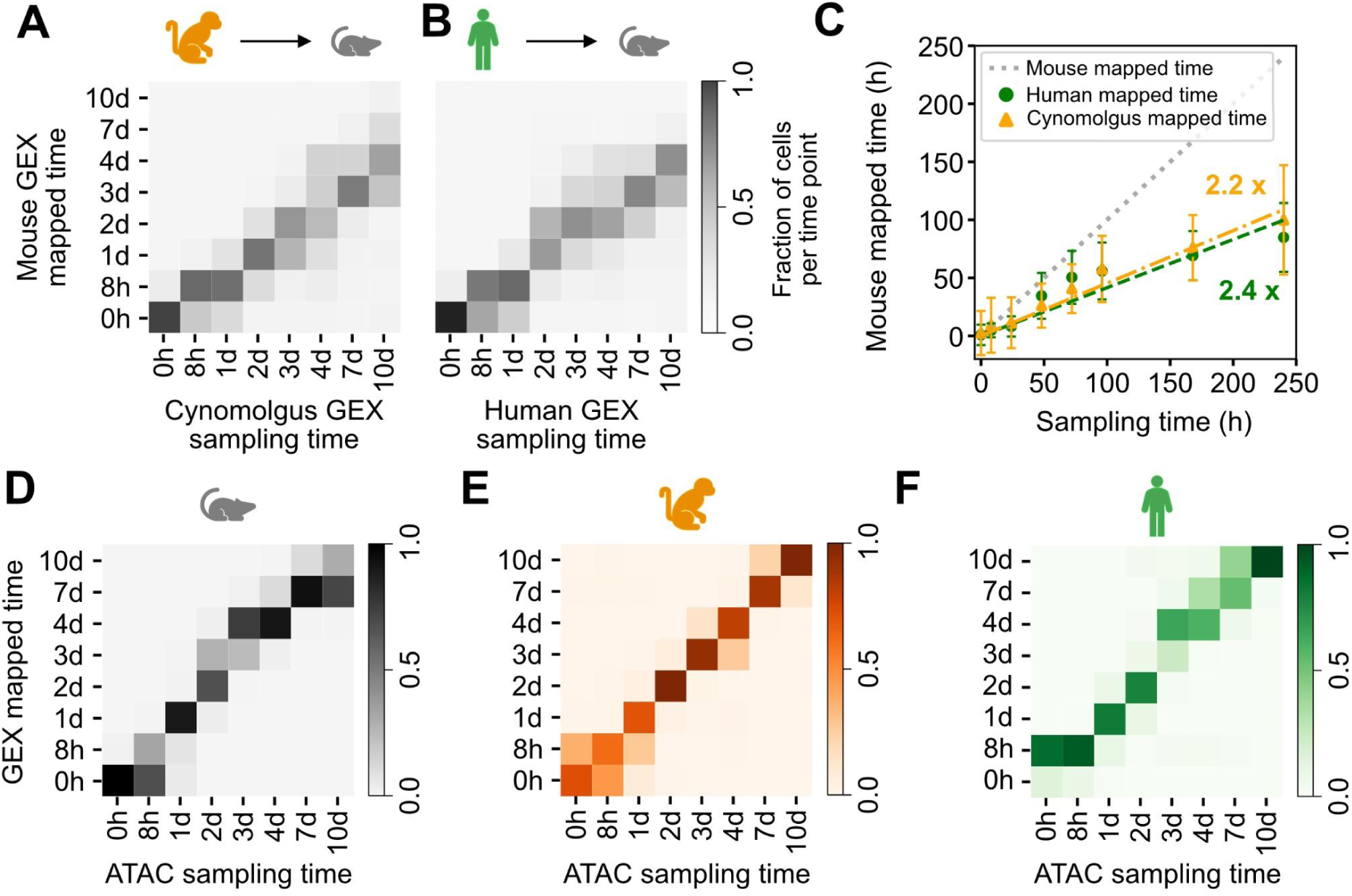
Inter-species single-cell mapping reveals that mouse cells differentiate 2.2 times faster than cynomolgus cells and 2.4 times faster than human cells. A-B. Heatmaps showing fractions of cynomolgus (A) and human (B) cells for each sampling time mapping to reference time points in mouse for scRNA-seq (GEX) data. Cells in cynomolgus and human differentiate significantly slower, e.g., most of the single cells sampled at day 10 in cynomolgus and human (last column in both matrices) are most similar, and thus mapped to mouse cells at day 4. C. Linear regression of mapped scRNA-seq data. Data points show the mean mapped time of human (green circles) and cynomolgus (orange triangles) cells on the mouse reference data. Lines are linear fits to the data for human (green, y = 0.45x, R^2^ = 0.95) and cynomolgus (orange, y = 0.42x, R^2^ = 0.84), indicating that mouse differentiation is 2.2 and 2.4 times faster than cynomolgus and human, respectively. Error bars indicate standard deviation. Line of unity (gray) shown as reference. D-F. Heatmaps show high correlations between simulated gene expression data from scATAC-seq and gene expression determined by scRNA-seq in mouse (D), cynomolgus (E) and human (F).

To test if similar species-specific dynamics could be observed at the chromatin level, we applied a similar analytical approach to scATAC-seq data. Comparing the simulated gene expression from scATAC-seq data (see Methods for details) with scRNA-seq data showed high accordance for all three species (**Figures 2D-F**). Using this simulated gene expression data, we mapped human and cynomolgus onto the mouse UMAP embedding and correlated human and cynomolgus sampling time and mouse mapped time for scATAC-seq (**Figures S2C** and **S2D**). Using the same linear regression model, mouse cells differentiated 1.9 times faster than human cells and 1.7 times faster than cynomolgus cells (**Figure S2E**). These results demonstrate that species-specific differences in differentiation speed are reflected at the level of chromatin accessibility.

### Species-specific neural differentiation speeds can be uncoupled from cell growth and the cell cycle

Pluripotent cells from fast-differentiating species have shorter cell cycles than those from slow-differentiating species^4,6^. To test if differences in cell cycle length and structure can quantitatively account for the species-specific differentiation speeds, we generated pluripotent stem cell lines expressing the PIP-FUCCI sensor^21,22^. Live-cell imaging of these reporter lines (**Figures S3A** and **S3B**) showed that the total cell cycle duration differed in the harmonized pluripotency conditions (**Figures 3A and S3C**): Mouse cells exhibited the shortest cell cycle with an average duration of 10.1 ± 0.9 hours (mean ± se, N = 2, n ≥ 39 each), while the cell cycle was longest in human cells with an average of 14.8 ± 0.1 hours (N = 2, n ≥ 40 each), which corresponds to a 1.47-fold difference between mouse and human cells. Cynomolgus cells fell in between mouse and human cells with an average cell cycle duration of 14.3 ± 0.9 hours (N = 2, n ≥ 39 each), 1.42-fold the average mouse duration. Although this variation in total cell cycle corresponds qualitatively with differences in the timing of cell differentiation between the species, it does not quantitatively account for the > 2-fold difference in differentiation speed (**Figure 2**). Cells from all three species exhibited the characteristic cell cycle structure of pluripotent cells, with a stretched S phase and a very short G1 phase (**Figure 3A**). When normalized to total cell cycle length, the distribution of the different phases was similar between cells from the three species (**Figure S3D**), arguing against regulation of developmental timing through changes of cell cycle structure^23,24^.

**Figure 3:**
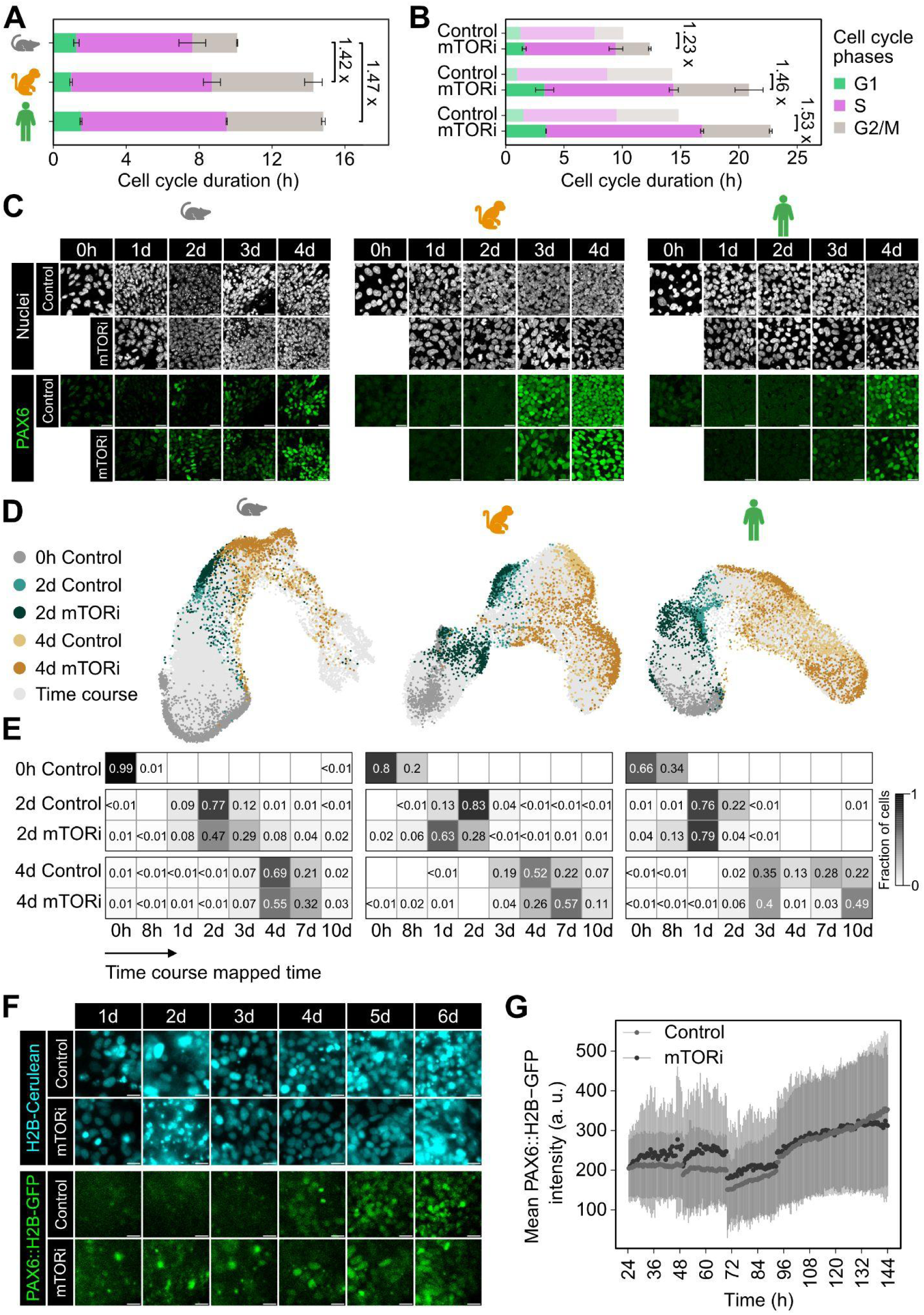
Changing cell cycle duration with mTOR inhibition does not affect differentiation speed. A. Bar graphs showing the mean duration of G1 (green), S (pink) and G2/M (gray) cell cycle phases in mEpiSCs, cyiPSCs, and hESCs measured by time-lapse imaging of the PIP-FUCCI reporter. Cynomolgus and human cells cycle approximately 1.42 times and 1.47 times slower than mouse cells, respectively, but cells from all three species have a similar distribution of G1, S, and G2M phases. Data from two independent experiments with n ≥ 39 cells each. Error bars indicate standard error. B. Measurements of cell cycle lengthening upon mTOR inhibition. Bar charts show mean G1 (green), S (pink) and G2/M (gray) cell cycle phase durations in cells treated with 50 nM INK128. Comparison with measurements in control cells (transparent, reproduced from A) reveals a 1.23-fold (mouse), 1.46-fold (cynomolgus) and 1.53-fold (human) lengthening of the cell cycle upon mTOR inhibition. Data from two independent experiments with n ≥ 40 cells each. Error bars indicate standard error. C. Differentiation time course of mouse (left), cynomolgus (middle) and human cells (right) fixed after 0 hours and 1, 2, 3, and 4 days of NPC differentiation in the absence (control) or presence of mTOR inhibition, stained for the early neural marker PAX6 (green). mTORi has no discernable effect on PAX6 expression onset. Scale bars, 20 µm. D. UMAP projection of single cell transcriptomes of mTORi-treated (dark teal and brown) and control cells (light teal and brown) onto the time course reference data set (gray). mTORi-treated and control cells occupy similar regions in UMAP space when mapped onto the time-resolved reference data set. E. Heatmap showing the proportion of cells within a sample assigned to a specific time point of the reference data set. F. Daily stills from time-lapse imaging of PAX6::H2B-GFP; H2B-Cerulean reporter cells differentiated in absence or presence of INK128. The INK128 dose used in imaging experiments was slightly reduced to 40 nM to improve cell viability during live cell microscopy. Constitutively expressed H2B-Cerulean to indicate nuclei positions shown in cyan, PAX6::H2B-GFP shown in green. G. Quantification of experiment shown in F. Data points show mean nuclear PAX6 reporter intensity across all cells in each frame. Sudden drops in fluorescence intensities at around 48, 72, and 96 h are caused by media changes. Data from control cells in light gray, data from mTOR-treated cells in dark gray. Error bars indicate standard deviation.

To functionally test how strongly differentiation speed is coupled to cell proliferation, we slowed down cellular growth by targeting mTOR with the ATP site inhibitor INK128 (mTORi). Partial inhibition with 50 nM of mTORi led to a species-specific extension of cell cycle durations: Mouse was affected least (1.23-fold extension compared to control) and human the most (1.53-fold), with cynomolgus in between (1.46-fold) (**Figure 3B**). If differentiation speed depended on growth and proliferation, the expression onset of neural markers would be expected to take approximately 50% longer in primate cells under mTOR inhibition compared to control cells. However, immunofluorescence of a differentiation time course did not reveal a strong delay in PAX6 (**Figure 3C**) and SOX1 onset (**Figure S3E**) upon mTOR inhibition in any of the species. Only OCT4 downregulation appeared to be slightly slower under mTOR inhibition in human cells, which could be caused by reduced protein dilution as a result of less frequent cell divisions (**Figure S3F**). To globally assess the differentiation status of mTOR-inhibited cells, we performed scRNA-seq of mouse, cynomolgus and human cells at 0 hours (pluripotent control) and days 2 and 4 of differentiation with and without mTORi (**Figures S3G-I**). To minimize batch effects, we multiplexed cells within each time-point (see Methods for details). Using Scanpy’s ingest tool^20^, we integrated the data from mTORi-treated and control cells with the time-resolved scRNA-seq differentiation time course to determine the reference time point to which untreated and mTOR inhibitor-treated cells best correspond to. In mouse, mTORi-treated cells mapped onto the same reference time point as untreated cells, showing that mTOR inhibition did not delay neural differentiation (**Figures 3D** and **3E**, left). In the primate species, day 2 mTOR-inhibited cells were slightly delayed compared to untreated cells, however, day 4 mTOR-inhibited cells preferentially mapped to later time points than untreated cells (**Figures 3D** and **3E**, middle and right). The overall tendency of both INK128-treated and untreated primate cells to map to earlier time points of the reference dataset than expected may be explained by different methods used to measure transcriptomes in the two experiments (see Methods for details). Still, the finding that cells map to similar reference time-points irrespective of INK128 treatment suggested that there is no systematic differentiation delay upon mTOR inhibition. To rule out the possibility that mTOR inhibition had more subtle effects on differentiation timing that might be missed due to the limited resolution of the reference dataset, we performed time-lapse imaging of a PAX6 reporter line^25^. We observed almost simultaneous onset of PAX6 reporter expression with and without mTOR inhibition (**Figures 3F** and **3G**). In sum, these findings show that differentiation can be uncoupled from growth, cell proliferation and cell cycling. Thus, species-specific neural differentiation speed must be determined by alternative mechanisms.

### Identification of candidate regulators of developmental speed through comparative cross-species analyses

We next applied a three-step analysis pipeline on our time course scRNA-seq dataset to identify biological mechanisms and specific candidate genes other than cell cycle that might account for species-specific differentiation speeds (**Figure 4A**). As a first step, we curated gene lists from GO categories such as energy metabolism, protein biosynthesis, and epigenetic regulation and modification that had previously been implied in developmental speed (**Table S1**)^4,5,13–15,26–30^. Normalizing the data for each biological pathway revealed unique expression profiles of these broad gene groups for each species: Human cells showed higher expression of epigenetic regulators and DNA modifiers, while cynomolgus cells had increased expression of genes associated with glycogen metabolism and biological oxidations. Mouse cells exhibited the highest upregulation in glucose-related metabolic processes. The tricarboxylic acid (TCA) cycle and nicotinamide adenine dinucleotide (NADH) metabolism displayed similar patterns across all species (**Figure 4B**).

**Figure 4:**
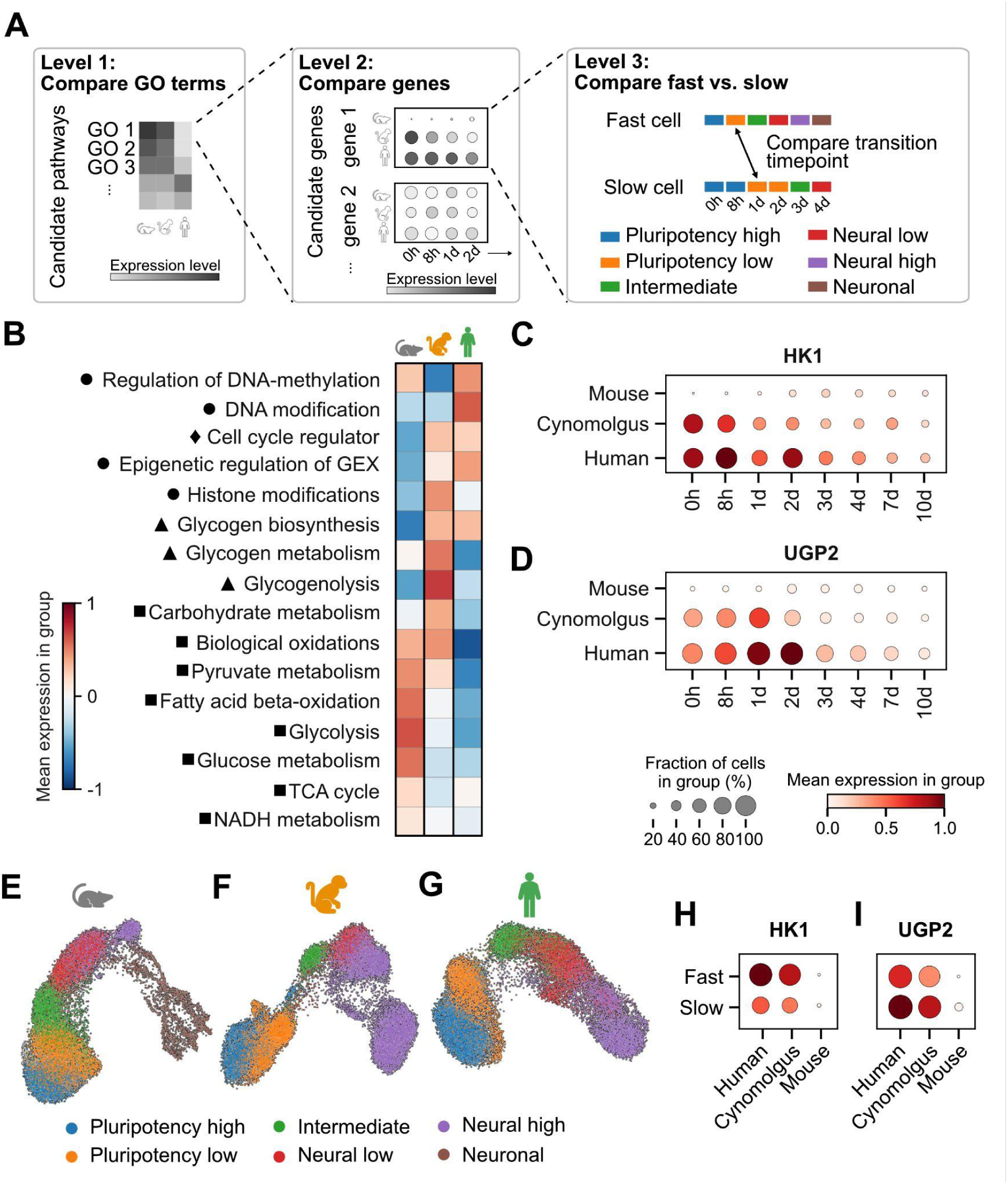
Glycogen biosynthesis related UGP2 is upregulated in slower species and in slower cells. A. Schematic overview of the approach used to identify candidate mechanisms and genes involved in differentiation speed. In the first level, candidate GO terms were compared across species. At the second level, GO terms of interest were further analyzed for individual genes that displayed distinct dynamics between species and time points. In the third level, these selected genes were compared between cells with fast and slow differentiation speed within each species. Here, cells were categorized into different clusters representing differentiation states (e.g., pluripotency high or low). For example, a cell sampled 8 hours post-induction categorized as pluripotency low was classified as fast-differentiating, whereas a cell sampled 1 day post-induction with the same state was labeled as slow-differentiating. B. Heatmap showing normalized mean expression of selected biological pathways across all three species. Circle marks epigenetic regulation, diamond cell cycle regulation, triangle glycogen related pathways and square metabolic pathways. C-D. Dots plots of HK1 (C) and UGP2 (D) expression levels during NPC differentiation in mouse, cynomolgus, and human. E-G. Progressive differentiation states for mouse (E), cynomolgus (F), and human (G) shown on the UMAP embeddings of single-cell gene expression data. Colors correspond to pluripotency high - blue; pluripotency low - orange; intermediate - green; neural low - red; neural high - pink; neuronal - brown. The neuronal cluster is only present in mouse cells (E), in line with their more extensive differentiation trajectory. H-I. Dot plots comparing HK1 (H) and UGP2 (I) expression levels in fast- and slow-differentiating pluripotency-low cells across the different species. UGP2 (I) expression levels are higher both in slowly differentiating species, as well as in slowly differentiating cells from the same species, whereas HK1 expression shows the reverse behavior (H).

In a second step, we asked which individual genes were responsible for the species-specific prevalence of selected biological processes. To address the possibility that such a species-specific preponderance was driven by differences in differentiation states between cells from the three species, we took into account the expression dynamics of individual genes across the differentiation time-course (**Figure 4A, middle)**. We specifically focused on the gene groups associated with the terms “glycogen biosynthesis” and “glycolysis”, because the former showed stronger average expression in the slow primate species, whereas genes in the latter group were on average more highly expressed in fast-differentiating mouse cells (**Figures S4A** and **S4B)**. Within each group, only a few genes showed strongly different expression between species and time points. Two such genes of interest were Hexokinase 1 (HK1) from the group “glycolysis”, which is responsible for the phosphorylation of glucose to glucose-6-phosphate during the initial step of glycolysis, and UDP-glucose pyrophosphorylase 2 (UGP2) from the group “glycogen biosynthesis”, which encodes an enzyme critical for re-routing glucose from glycolysis into glycogen storage. Both HK1 and UGP2 exhibited high expression in primate cells at the onset of neural differentiation, while remaining absent in mouse cells throughout the entire differentiation process (**Figures 4C** and **4D**).

In the third step of our analysis, we sought to leverage the heterogeneous differentiation of cells from the same species in our time course transcriptomic data to devise an independent test for the involvement of a candidate gene in controlling differentiation speed. We hypothesized that if a candidate gene affected differentiation speed, we would find differences in its expression levels when comparing slow- and fast-differentiating cells from the same species (**Figure 4A, right**). To distinguish fast- and slow-differentiating single cells, we first employed unsupervised Leiden clustering of the whole dataset for each species, to define six specific stages of differentiation: pluripotency high, pluripotency low, intermediate, neural low, neural high, and neuronal (**Figures 4E-G**). We then defined sets of marker genes for each cluster (see Methods for details). These genes showed a similar expression sequence across species, albeit with different dynamics (**Figures S4C-E**), such that mouse cells reached the same differentiation stage faster than primate cells (**Figures S4F-H**). Based on the marker gene sets, we calculated gene expression scores for each cell and each cluster (**Figures S4I-K**), and applied a stringent threshold of this score to select cells with highly similar gene expression profiles that closely matched the characteristic profile of the respective cluster. Finally, we compared the expression of previously identified genes of interest in cells that reached a high marker gene score at early time points (fast-differentiating cells) to their expression in cells that reached the same marker gene score at later time points (slow-differentiating cells). This analysis revealed that UGP2, but not HK1, consistently exhibited higher expression in slow-differentiating cells compared to fast-differentiating cells (**Figure 4H** and **4I**). This effect was observed across all species and was particularly pronounced in cynomolgus and human cells, likely due to their overall higher UGP2 expression levels compared to mouse cells. The higher expression of UGP2 in slow-differentiating cells, both in intra- as well as cross-species comparisons, make it a strong candidate for the control of cell differentiation speed.

### UGP2 expression regulates species-specific glycogen storage and neural differentiation dynamics

Finally, we tested the functional roles of species-specific UGP2 expression for cellular carbohydrate metabolism and neural differentiation. Consistent with RNA levels, similar amounts of UGP2 protein were detected in human and cynomolgus wild-type pluripotent stem cells, but UGP2 protein was nearly absent in mEpiSCs (**Figure 5A**). UGP2 protein was also undetectable in a previously described human ESC UGP2 knockout (KO) line^31^ (**Figure 5A**). Species-specific UGP2 protein levels translated into glycogen storage levels: While human pluripotent stem cells stored high levels of glycogen, cynomolgus and mouse cells only contained approximately 50% and 10% of the per cell glycogen levels detected in human cells, respectively (**Figure 5B**). These differences in glycogen levels are reminiscent of the trends in differentiation speeds across the species, and establish that glycogen storage levels as a new species-specific cellular property. Glycogen levels were strongly decreased in UGP2 KO hESCs (**Figure 5B**), demonstrating that glycogen storage depends on, and is likely regulated through UGP2 expression. We then asked if loss of UGP2 would alter differentiation dynamics. In human UGP2 KO cells, RT-qPCR revealed a consistent upregulation of neural markers SOX1, PAX6, and FOXG1 compared to wild-type cells at several stages (**Figure S5A**). When staining for the late-stage neural marker FOXG1 in a differentiation time course, we found the first FOXG1-expressing cells by day 9 in UGP2 KO cells, but only at day 14 in the wild-type, further indicating accelerated neural differentiation upon loss of UGP2 (**Figures 5C** and **5D**). To assess changes in differentiation speed upon loss of UGP2 transcriptome-wide, we performed scRNA-seq of the UGP2 KO cells in pluripotency conditions and after 7 days of neural differentiation (**Figure 5E**). Mapping mutant transcriptomes to the reference time course data revealed that human UGP2 KO cells at 0 hours resemble 8-hour and 0-hour reference cells, while day 7 UGP2 KO cells mapped mostly to day 10 wild-type cells, indicating faster neural differentiation upon loss of UGP2 (**Figures 5F** and **5G**). Gene enrichment analysis for differentially regulated genes in KO samples showed several neural-related pathways within the top 100 upregulated KEGG pathways, such as neuroactive ligand-receptor interaction and various synaptic pathways (**Figure 5H**). Interestingly, glycolysis/gluconeogenesis pathways were also represented in the top 100 enriched pathways, suggesting autoregulation at the transcriptional level. We also generated a UGP2 KO cyiPSC line to test if a similarly accelerated neural differentiation could be observed in this species. UGP2 protein was undetectable in UGP2 KO cyiPSCs (**Figure S5B**), and *FOXG1* mRNA was more highly expressed compared to wild-type cells from day 4 onwards (**Figure S5C**). In an scRNA-seq experiment however (**Figure S5D**), mapping mutant transcriptomes onto the reference time-course data from Figure 1 did not reveal major changes to differentiation speed upon loss of UGP2 in cynomolgus cells (**Figures S5E** and **S5F**). This smaller effect of the loss of UGP2 on differentiation dynamics in cynomolgus compared to human cells might be related to overall lower glycogen storage levels in cynomolgus cells, and additionally be obscured by the low time resolution of the reference dataset at the later time points. Still, these experiments consistently demonstrated accelerated neural differentiation in human cells lacking functional UGP2, suggesting that UGP2 activity in wild-type cells slows down the pace of neural differentiation.

**Figure 5:**
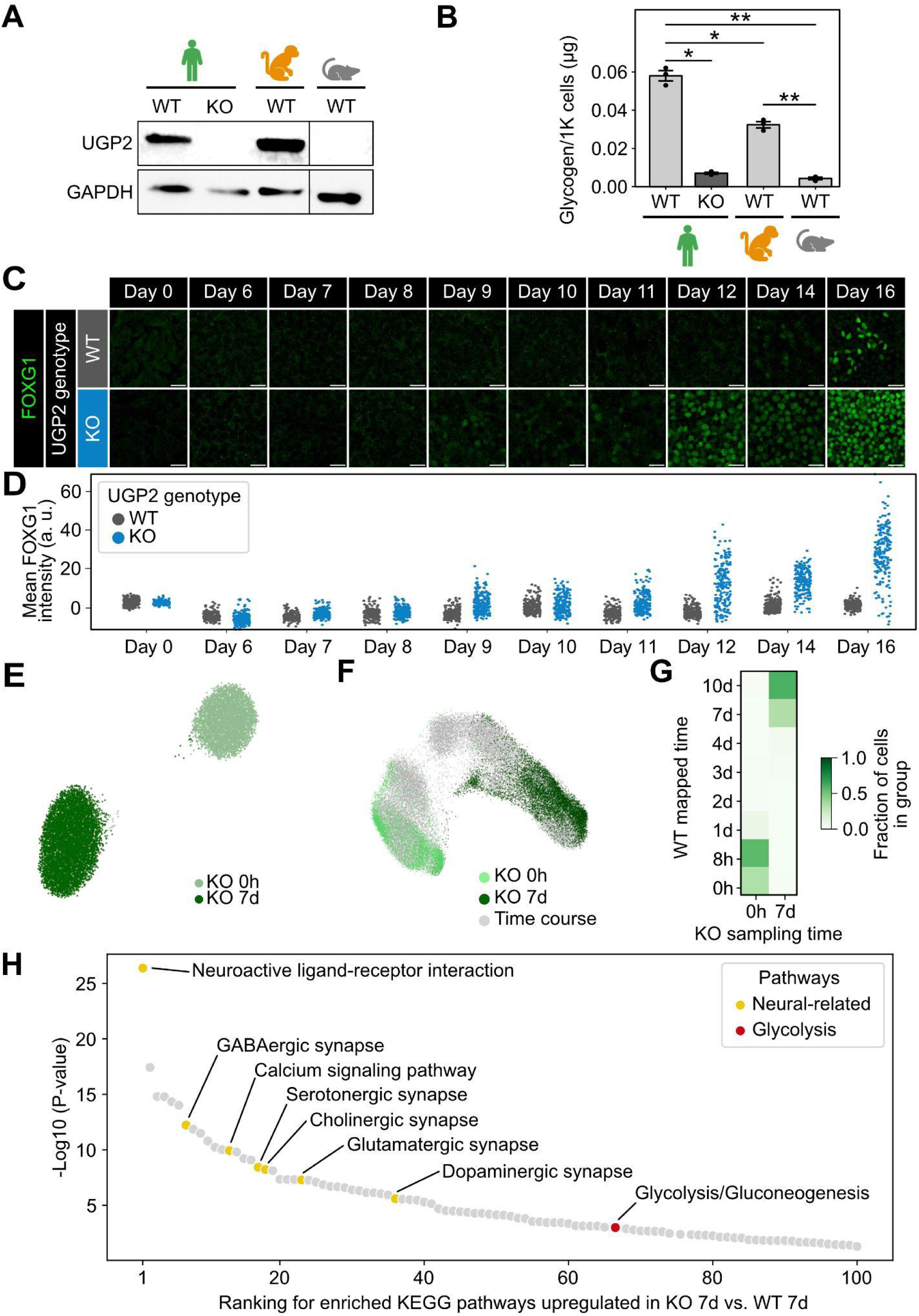
Loss of UGP2 accelerates human NPC differentiation. A. UGP2 protein expression in wild-type (WT) cells from the three species, and in human UGP2 KO cells, detected by Western blotting. Human and cynomolgus cells have similar UGP2 expression levels, but UGP2 is absent in human UGP2 KO and mouse wild-type cells. GAPDH is used as housekeeping control. B. Mean glycogen content in pluripotent cells from the three species. Glycogen levels are lowest in mouse, and 7.5 and 13.5 times higher in cynomolgus and human cells, respectively. Points indicate data from N = 3 independent experiments, error bars indicate standard error. ns: adjusted p-value * p < 0.05, ** p < 0.01, as determined by an unpaired Student’s t-test with Bonferroni correction. C. NPC differentiation time course of human UGP2 wild-type and KO cells fixed and stained for the anterior neural/forebrain differentiation marker FOXG1 (green). FOXG1 expression is detected earlier in KO than in wild-type cells. Scale bars, 20 µm. D. Quantification of immunostaining shown in (C). Single cell FOXG1 expression levels were measured in nuclear masks determined by segmentation in the Hoechst33258 channel via StarDist 2D using the Versatile Fluorescent Nuclei model^32^. For visualization, the number of cells was reduced to a maximum of 200 cells per sample. A background value measured in a 200×200 μm region of interest of each image was subtracted and outliers defined in each image as all values outside a range of 3 times the interquartile range below or above the first and third quartile respectively were removed in each channel. Only cells with an area ≥ 10 μm^2^ were considered. E. UMAP representation of scRNA-seq data of human UGP2 KO cells before (0 hours) and after 7 days of NPC differentiation. F. UMAP projection of single-cell transcriptomes of UGP2 KO cells at 0 hours (light green) and 7 days (dark green) of differentiation onto the time-resolved reference data set (gray). G. Heatmap showing the fraction of UGP2 KO cells at 0 hours and 7 days of NPC differentiation that map to specific time points of the wild-type reference data set. Day 7 UGP2 KO cells mostly map to wild-type day 10 and are hence developmentally advanced compared to the wild-type. H. KEGG pathway enrichment using differentially expressed genes in UGP2 KO versus wild-type cells at day 7 of NPC differentiation. Top 100 picks sorted by p-value are displayed, neural related pathways are highlighted in yellow, glycolysis in red. Neural related pathways are upregulated in the UGP2 KO.

## Discussion

Here we asked why differentiation speeds of pluripotent cells from different species are so different, despite similar genetic makeup and growth in the same media. Using single-cell multiomic data from neural differentiation in a harmonized three-species stem cell panel, we determined precise scaling factors of differentiation speed along the neural lineage speed between mouse, cynomolgus and human cells. We showed that differentiation speed differences are reflected in chromatin dynamics, but can be uncoupled from cellular growth and cell cycling. Leveraging the multi-species and single-cell nature of our data, we identified several metabolic pathways as candidate regulators of developmental speed, and functionally tested the glycogen-storage regulator UGP2 as the first metabolic enzyme for the species-specific and cell-intrinsic control of differentiation speed.

The differentiation of cortical neural progenitors has been one of the first differentiation paradigms to study the cell-intrinsic control of differentiation speed with stem cell models^3^. Subsequently, the regulation of differentiation speed has been studied with protocols leading to several other cell identities, most prominently motor neurons and presomitic mesoderm^4–6^. The advantage of the neural progenitor differentiation approach is that it exploits a default cellular differentiation path in the absence of any extracellular signals, and therefore most cleanly reports on the cell-intrinsic basis of developmental timing. Still, the 2.4-fold faster differentiation of mouse compared to human cells measured by us closely aligns with previously reported values of 2.5 for motor neuron differentiation^6^ and 2.6 for segmentation clock oscillations^5^. These similarities suggest that the scaling of cell-intrinsic differentiation speeds is largely conserved across different protocols, labs and experimental paradigms.

In contrast to the strong speed differences between mouse and human cells, we found that cynomolgus cells differentiate only marginally faster than human cells. Using segmentation clock oscillations *in vitro* as a model, Lázaro et al. have recently suggested that species-specific embryogenesis length is most predictive for cell differentiation speeds *in vitro*^13^. Our data supports this notion: Despite an almost two-fold difference in total gestation times, the end of embryogenesis at Carnegie stage 23 is reached after similar times in monkeys (46 days)^33^ and humans (58 days)^34^.

By measuring differentiation at different levels of regulation, we demonstrated that species-specific chromatin dynamics are closely aligned with transcriptome changes. Consistent with this finding, the slow maturation of human cortical neurons has been linked to an epigenetic barrier that could be overcome by the targeted inhibition of modifiers of chromatin accessibility^35,36^. Together, these results therefore enforce the concept that, at least in neural differentiation, epigenetics and chromatin dynamics are a key determinant of developmental speed.

An array of cell physiological parameters has been implicated in the control of developmental speed, such as mitochondrial activity^4,15^, protein half-life^6^ and general biochemical reaction rates^5^. Our experiments with the mTOR inhibitor INK128 suggest that previously proposed links between translation rate and differentiation speed may not apply to neural differentiation. The significantly lengthened cell cycle upon mTOR inhibition indicates that we have succeeded in slowing down anabolic processes, as would be expected by targeting mTOR as a central hub of metabolism that promotes cell growth through the upregulation of protein translation and ribosome biogenesis^37^. Still, we could not detect any differences in the onset of neural marker gene expression between untreated and mTOR inhibited cells, suggesting that the progression of the neural differentiation programme proceeds independently of general anabolic processes and the cell cycle. Although it remains a possibility that mTOR inhibition has pleiotropic effects on both neural differentiation cell growth that are independent of each other and could compensate^38^, our *in vitro* results align with findings from an *in vivo* study, which likewise demonstrated that neural progenitor differentiation progressed through successive stages irrespective of genetic interventions to the cell cycle^39^. Previously reported effects of translation rate downstream of mitochondrial activity on differentiation speed^4^ may therefore act through specific pathways rather than through the global reduction of anabolic rate, or only apply in specific differentiation paradigms.

Exploiting the single-cell multi-species nature of our time course transcriptomic dataset, we searched for candidate pathways and genes for developmental speed control. Focusing on UGP2 as one prominent hit from this analysis, we have identified glycogen storage as a new species-specific trait of pluripotent cells that are otherwise in comparable developmental states. Besides generating precursors for glycogen synthesis and thereby energy storage, UGP2 is also involved in the synthesis of extracellular matrix components such as hyaluronan, and its product UDP-glucose is a substrate for protein glycosylation^40^. Which of the diverse functions of UGP2 underlies the accelerated neural differentiation in the UGP2 knockout cells remains at this point unclear. It is tempting to speculate that increased energy storage in the form of glycogen is functionally related to an overall reduced metabolic activity and subsequent slower differentiation speed in primate compared to mouse cells. Such a relationship has been demonstrated in flatworms, where an increase in energy storage in cells from larger organisms results in a slower metabolic rate compared to cells from smaller organisms^41^. Intriguingly, flatworm cells with a lower per-mass metabolic rate are bigger than metabolically more active cells, just as human pluripotent cells are bigger than mouse pluripotent cells^4^. It is therefore an attractive possibility that the partitioning of high-energy metabolites between storage and catabolism is a general strategy to tune developmental speed in evolution.

Overall, our study adds glucose metabolism and glycogen storage to the growing list of cell-intrinsic mechanisms that control differentiation speed. Given the diversity of mechanisms that has been reported so far, it seems likely that each developmental lineage may rely on a slightly different combination of mechanisms to tune its differentiation speed. Understanding how scaling factors arise from diverse combinations of mechanisms is an important question for future research.

## Methods

### Cell lines and culture conditions

H9 human embryonic stem cells (ESC; WiCELL Research Institute), 129S2C1a mouse epiblast stem cells (EpiSC)^42^ and 56A1 cynomolgus induced pluripotent stem cells (iPSC)^43,44^ were plated on Matrigel (Corning) diluted 1:100 in DMEM/F12 (PAN Biotech) and gradually adapted to UPPS^19^ media consisting of StemMACS iPS-Brew XF (Miltenyi Biotec) supplemented with 1 µM IWR-1 (Sigma Aldrich) and 0.5 µM CHIR (R&D Systems) post passaging over four days. Afterwards, culture media was exchanged daily and cells were passaged with 0.5 mM EDTA (Promega) every two to four days as clumps. Cell identity was verified through the alignment of scRNA-seq reads to the genomes of their respective species. Experiments involving human stem cells were performed in accordance with permissions obtained from the Robert-Koch-Institute (AZ: 3.04.02/0151 to MD to and AZ 3.04.02/0172 to CS). All reagents and resources are listed in **Table S2**.

### In vitro neural progenitor cell differentiation

PSCs were singularized using Accutase (Sigma Aldrich) and counted on the Countess II FL Automated Cell Counter (Invitrogen). For hESCs and cyiPSCs 1.25 x 10^5^ cells/cm^2^ and for mEpiSCs 6.25 x 10^4^ cells/cm^2^ were seeded onto Matrigel coated plates in UPPS media supplemented with 10 µM Y-27632 (R&D Systems) and incubated overnight, resulting in confluent cultures the following day for each species. For neural induction, media was exchanged to NPC differentiation and maintenance media consisting of 1:1 DMEM/F12 and Neuropan (PAN Biotech) supplemented with 0.5x N2 and B27 supplements, 1x MEM-NEAA, 1x GlutaMAX, 0.1 mM 2-Mercaptoethanol, 5 µg/mL Insulin, human recombinant (all Thermo Fisher Scientific), 10 µM SB431542 and 100 nM LDN193189 (both Peprotech). Differentiation was performed for maximal 14 days. Cells were washed with Dulbecco’s phosphate buffered saline (DPBS; Thermo Fisher Scientific) before daily media exchange.

### Quantitative reverse transcription PCR (RT-qPCR)

Cells were lysed and RNA was isolated using the RNeasy Mini Kit (Qiagen) following the manufacturer’s protocol. Subsequently, cDNA was synthesized from 500 ng RNA using the Verso cDNA Synthesis Kit (Thermo Fisher Scientific). For RT-qPCR, SYBR Green PCR Master Mix (Thermo Fisher Scientific), cDNA and 5 µM of each forward and reverse primer were combined to a total reaction volume of 10 µL. RT-qPCR was performed in 384-well-plates on a QuantStudio 12K Flex qPCR machine (Thermo Fisher Scientific) in technical and biological triplicates. Relative normalized expressions were calculated using the ΔΔC_T_ method. Primers are listed in **Table S3**.

### Flow cytometry

First, cells were dissociated to single cells using Accutase and collected. For surface marker staining, cells were once washed with FACS buffer (0.5% BSA and 2 mM EDTA in DPBS) followed by centrifugation at 300 rcf for 5 minutes and removal of the supernatant. Cells were incubated with primary antibodies for 30 minutes on ice. Then, after another centrifugation and removal of supernatant, cells were incubated with secondary antibodies for 30 minutes on ice, then washed and resuspended in FACS buffer. For intracellular staining, the Inside Stain Kit (Miltenyi Biotec) was used according to the manufacturer’s protocol. Cells were incubated with primary antibodies for 1 hour at room temperature and with secondary antibodies for 30 min on ice. After an additional wash with Inside Perm solution, cells were resuspended in FACS buffer for further analysis on a BD FACSAria III cell sorter (BD Biosciences). Flow cytometry data was analyzed using the FlowJo software. Antibodies and dilutions are listed in **Table S4**.

### Immunostainings

Cells were grown on chambered polymer coverslips (ibidi) and fixed for 15 min in 4% formaldehyde (ROTI Histofix). Before antibody incubation, fixed cells were rinsed and washed three times for 15 min in PBT-BSA (PBS with Ca^2+^ and Mg^2+^ containing 1% BSA and 0.1% Triton). Primary antibodies were added in PBT-BSA overnight at 4 °C and washed off three times for 15 min. Secondary antibodies were incubated for 2 h at room temperature and washed off three times for 15 min in PBT-BSA and twice 10 min in PBS. Imaging was performed in mounting medium (80% glycerol, 20% H_2_O, 4% w/v n-propyl-gallate). Antibodies and dilutions are listed in **Table S4**.

### PIP-FUCCI measurements

For cell cycle characterizations, we used the PIP-FUCCI sensor by Grant et al.^21^ (Addgene plasmid #118621) and amplified it with overlaps for a piggyBAC vector^45^ containing a CAG promoter and a puromycin resistance cassette. The piggyBAC vector was opened via NotI-HF and XhoI digest and the PIP-FUCCI construct cloned between the restriction sites using NEB HiFi Assembly. Mouse and human cells were lipofected using lipofectamine2000, and cynomolgus cells were nucleofected using the Neon transfection system with the piggyBAC PIP-FUCCI construct and a pBASE plasmid for transposition^45^ and subsequently kept under 1.5 µg/ml puromycin selection.

PIP-FUCCI imaging was performed in chimeric cultures consisting of PIP-FUCCI sensor cells and unlabeled, parental cells in ratios ranging from 1:10 to 1:100. This enabled us to track cells even in dense colonies and during NPC differentiation when cells are seeded at high densities. Cells were seeded a day prior to acquisition start as single cells in UPPS medium supplemented with 10 µM ROCK inhibitor. Primate cells were typically seeded at 80,000 cells/cm^2^ and mouse cells at 40,000 cells/cm^2^. Immediately before imaging, cells were washed and transferred to the desired media condition, typically UPPS +/- 50 nM INK128. Images were acquired in 10 min intervals over a course of 48 hours on an Olympus IX81 widefield microscope with a 20X objective (NA 0.75). Cells were kept in a humidified stage top incubator (ibidi) at 37 °C and 5% CO_2_ during image acquisition. Tracking was performed in Fiji (ImageJ v2.9.0) with TrackMate v7.9.2^46^ using the “Manual Tracking” option. Here, a circular region of interest (ROI) with a radius of 2.24 px was manually placed in the center of a nucleus in each frame of a track. mVenus and mCherry fluorescence were measured inside the ROI. Each track started at the first frame after cell division and ended with the last frame before the next division, so the total cell cycle duration was calculated as *end frame – start frame*. Cell cycle phase durations were calculated based on mVenus fluorescence. The G1/S transition was defined as the frame in which mVenus signal in the first half of the track (fluorescence intensity scaled between 1 and 2) after the first peak was closest to its half maximum (PIP_half-max_). The S/G2M transition was the frame in which mVenus signal had an increase of > 0 and kept rising at an average of ≥ 1.5% over the following five frames (PIP_rise_) in the second half of the track (scaled between 1 and 2). Thus, phase durations were calculated as follows:

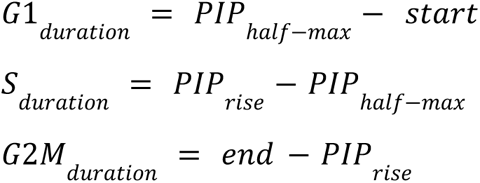

### PAX6 reporter time-lapse imaging

Human ESCs carrying a PAX6::H2B-GFP reporter^25^ were transfected with a piggyBAC vector containing an H2B-Cerulean-IRES-bsr construct under control of a CAG promoter to add a nuclear label for cell tracking^47^. After selection under 15 µg/ml blasticidin, polyclonal cells were seeded onto chambered polymer slides (ibidi) at a density of 250,000 cells/cm^2^ and grown in UPPS + 10 µM ROCK inhibitor overnight. NPC differentiation was induced at day 0, either without or with 40 nM INK128. After one day of NPC differentiation, image acquisition was started on an Olympus IX81 widefield microscope with an iXon 888 EM-CCD camera (Andor) and LED illumination (pE4000, CoolLED) on a 20X objective. All hardware components were controlled by MicroManager 2.0^48^. Images were taken in 30 min intervals for up to five days. Cells were washed daily and medium was changed. Time-lapse movies were segmented based on the nuclear H2B-Cerulean signal using StarDist 2D and the *Versatile (fluorescent nuclei) model*^32^ and the probability threshold set to 0.7 with otherwise default parameters. Accumulation of dead cells during differentiation and debris removal by daily washing led to artificial shifts in reporter signal intensity. To reduce this artifact, a Gaussian blur (radius = 50 µm) of the PAX6::H2B-GFP channel in each frame was generated and subtracted from the original image. PAX6::H2B-GFP intensity was measured on the resulting corrected image. Nuclei with an area ≤ 50 μm^2^ were filtered out and the frame average of PAX6::H2B-GFP intensity was calculated across all remaining nuclei per condition.

### CRISPR/Cas9 gene editing for the generation of cynomolgus iPSC UGP2 KO lines

CRISPR editing of cynomolgus iPSCs was carried out using a plasmid-free method, employing TrueCut Cas9 protein and TrueGuide sgRNAs (Invitrogen). The guide RNAs, designed to target the UGP2 gene locus in *Macaca fascicularis*, were generated using the Synthego CRISPR design tool and are detailed in **Table S3**. To generate KO cell lines, ribonucleoprotein (RNP) complexes were created by combining individual guide RNAs with TrueCut Cas9 protein, followed by a 10-minute incubation at room temperature and subsequent storage on ice for later use. Subsequently, 5 x 10^5^ cells were suspended in 100 µl of P3 Nucleofector™ solution (Lonza) per reaction, following the manufacturer’s protocol, and the prepared RNP complexes were added as required. The nucleofection process was carried out using the DN100 program on the Amaxa 4-D Nucleofector (Lonza).

After transfection, the cells were seeded onto 6-well plates coated with a 1:100 Matrigel in UPPS media supplemented with 10 µM ROCK inhibitor Y-27632 and incubated for 24 hours. Upon reaching 80% confluency, the cells were dissociated into single cells using Accutase as previously described. Subsequently, 10,000 single cells were plated onto 10 cm polystyrene dishes coated with a 1:100 Matrigel in UPPS media containing 10 µM ROCK inhibitor Y-27632 for 24 hours. The culture was maintained until visible colonies formed, and individual colonies were selected and transferred to 12-well plates, where they were cultured in UPPS media for further analysis and evaluation.

### Genomic DNA isolation, PCR, PCR clean-up and gel electrophoresis

To confirm successful homozygous mutagenesis of the UGP2 locus, genomic DNA was isolated from KO clones following the nucleofection of RNPs, utilizing the QIAamp DNA Mini Kit (Qiagen) and adhering to the manufacturer’s instructions.

PCR amplification was performed using DreamTaq Green DNA Polymerase (5 U/μL, Thermo Fisher Scientific), following the manufacturer’s protocol. Subsequently, the resulting PCR product was extracted using the QIAquick PCR purification kit (Qiagen). The PCR products were then examined using 1% agarose (Biozym) gels in 1x TAE buffer composed of 40 mM TRIS base, 20 mM acetic acid, and 1 mM EDTA (all Carl Roth). SYBR Safe DNA Gel Stain (Thermo Fisher Scientific) was incorporated into the gel at a dilution of 1:10,000. Electrophoresis was carried out at 100 V in TAE buffer. Subsequently, the DNA fragments were visualized using the ChemiDoc MP Imaging System (BioRad).

### Sanger sequencing

The PCR fragments were sent to Eurofins Scientific for Sanger Sequencing. Subsequently, the resulting sequences were analyzed utilizing SnapGene viewer software v7.1.1 to verify the accuracy of the intended deletions.

### Western blot

To evaluate the successful generation of KO cell lines at the protein level, Western Blot analysis was conducted. Total protein was extracted using RIPA buffer (Thermo Fisher Scientific) supplemented with 1x protease inhibitor (Roche). The lysed cells were then cooled on ice for 5 minutes, followed by centrifugation at 14,000 x g for 15 minutes. The resulting supernatant was transferred to a fresh tube. To measure protein concentration, Pierce™ BCA Protein Assay Kit was used according to the manufacturer’s instructions and 36 µg of total protein were used for subsequent analysis. 2x Laemmli buffer (BioRad) was added to all samples and incubated at 95°C for 5 minutes. The prepared protein sample was applied to a 7.5% Mini PROTEAN® TGX stain-free gel (BioRad) for electrophoresis, using SDS running buffer containing 1x Tris/glycine (BioRad) and 3.5 mM SDS (Serva Electrophoresis). Wet blotting onto a nitrocellulose membrane was performed for 1 hour at 100 V using blotting buffer with 1x Tris/glycine and 20% methanol (Serva Electrophoresis).

Subsequently, the membrane was blocked for 1 hour at room temperature in TBST buffer, consisting of 20 mM TRIS base, 150 mM NaCl, 0.1% Tween-20, and 5% milk powder (all Carl Roth). The primary antibody was diluted 1:1000 in blocking buffer, applied to the membrane, and incubated overnight at 4°C. Antibodies used for Western blot are listed in **Table S4**. After three washes with TBST buffer, the membrane was incubated with a secondary antibody conjugated with horseradish peroxidase (Sigma Aldrich) at a 1:10,000 dilution in blocking buffer for 2 hours at room temperature. Following another three washes of 15 minutes each in TBST buffer, the membrane was stained with Clarity Western ECL Substrate and visualized after a 2-minute incubation using the ChemiDoc MP Imaging System (BioRad).

### Glycogen measurements

Glycogen contents were measured using the Glycogen Assay Kit ab65620 (Abcam) following manufacturer’s instructions. Cells were harvested, washed with PBS and resuspended in cold H_2_O. For glycogen content normalization, cells were counted using a Countess Automated Cell Counter (Invitrogen). Enzymatic inactivation was performed at 100°C for 10 min and glycogen was hydrolysed to glucose and detected using the OxiRed probe following the manufacturer’s instructions. All samples were measured in technical duplicates and a sample background control where glycogen is not hydrolysed to glucose was performed. A standard curve with glycogen concentrations ranging from 0 (blank) to 0.2 µg glycogen/well was used to calculate glycogen contents. All samples, sample background and standard wells were measured at 535/587 nm (Ex/Em) on a Tecan Plate Reader. Duplicate measurements were averaged and sample background readings as well as a blank value were subtracted from sample readings. The standard readings were plotted against the glycogen concentration/well, and a linear fit was performed to calculate glycogen contents in each sample well. The glycogen content [µg/µl supernatant] was then normalized to the number of cells in solution before homogenization.

### Single-cell sequencing experiments and analysis

#### Nuclei isolation for single-cell multiome sequencing

Nuclei isolation was only performed for time course single-cell multiome sequencing samples. At the determined time points during NPC differentiation, hESCs, mEpiSCs, and cyiPSCs were individually dissociated to single cells using Accutase. Subsequently, 2.5 x 10^5^ cells per species were collected in a 2 mL DNA LoBind Tube (Eppendorf) and combined to a single sample for further processing. Nuclei isolation was performed according to the demonstrated protocol provided by 10x Genomics with some modifications. Briefly, cells were centrifuged at 300 rcf for five minutes and the supernatant was removed. The cell pellet was gently resuspended in 100 µL lysis buffer (10 mM Tris-HCl pH 7.4, 10 mM NaCl, 3 mM MgCl_2_ (all Sigma Aldrich), 0.05% Tween-20 (Carl Roth), 0.05% NP40 (Sigma Aldrich), 0.01% Digitonin, 1% BSA, 1 mM DTT and 1 U/µL RNase inhibitor (all Thermo Fisher Scientific)) and incubated for three minutes on ice. After incubation, the sample was washed by the addition of 1 mL wash buffer (10 mM Tris-HCl pH 7.4, 10 mM NaCl, 3 mM MgCl_2_, 0.1% Tween-20, 1% BSA, 1 mM DTT and 1 U/µL RNase inhibitor), followed by centrifugation at 1000 rcf for 5 minutes and removal of the supernatant. Washing was repeated two more times. After the last washing step, the nuclei pellet was resuspended in Diluted Nuclei buffer (1x Nuclei Buffer (10x Genomics), 1 mM DTT and 1 U/µL RNase inhibitor). The concentration was determined and the sample was checked for successful nuclei isolation by visual examination under the microscope using Trypan Blue (Invitrogen). While the following library preparation was performed on single isolated nuclei, the term ‘single-cell sequencing’ is employed for the time course multiome data.

#### Library preparation and sequencing

Time course scATAC and scRNA libraries were prepared using the Chromium Next GEM Single Cell Multiome ATAC + Gene Expression Reagent Bundle (10x Genomics) according to the manufacturer’s protocol. Samples were collected at 0h, 8h, 1d, 2d, 3d, 4d, 7d and 10d post neural induction and cells of all three species were combined in equal amounts prior to the library preparation. The aimed target recovery was 10,000 single nuclei for sequencing. Libraries were sequenced on an Illumina NovaSeq 6000.

For scRNA-seq in the mTOR inhibition experiments, the Chromium Next GEM Single Cells 3’ Reagent Kits v3.1 (Dual Index) with Feature Barcode technology for Cell Multiplexing (10x Genomics) were used following the manufacturer’s instructions with minor modifications. Neural differentiation was initiated as described, and 50 nM INK128 was added to half of the cells. Cells were harvested at 0h, 2d, and 4d of NPC differentiation. Labeled samples were pooled and loaded onto a Chromium Next GEM Chip, with target recoveries of 12,000 cells for day 0 and 22,000 cells for days 2 and 4. Sequencing reads were obtained through NovaSeq X Plus PE150.

For the UGP2 KO experiments, scRNA-seq was performed using the Chromium Next GEM Single Cells 3’ Reagent Kit v3.1 (Dual Index) according to the manufacturer’s instructions, with samples collected at 0h and 7d post neural induction. Equal amounts of cynomolgus and human cells were combined for a target recovery of 10,000 cells for subsequent library preparation and species assignment. Libraries were sequenced on an Illumina NovaSeq 6000.

#### Preprocessing

For time course single-cell multiome sequencing experiments, CellRanger ARC (v2.0.0) provided by 10x Genomics was used to demultiplex binary Illumina base call (BCL) files into FASTQ files, align the data against the GRCh38 (human), GRCm39 (mouse), and Macaca_fascicularis_6.0 (cynomolgus) reference, filter reads, as well as count barcodes and scRNA and scATAC molecules to generate feature-barcode matrices. Mapping of the mTOR inhibition experiments was performed using CellRanger 7.2.0 (10x Genomics). Gene Expression libraries were mapped against the GRCm38 (mouse), GRCh38 (human), and Macaca_fascicularis_6.0 (cynomolgus) genomes. Mapping of the scRNA-seq data of the UGP2 KO was performed using CellRanger 7.1.0 (10x Genomics) and reads were aligned against the GRCh38 (human), and Macaca_fascicularis_6.0 (cynomolgus) genomes.

Subsequent preprocessing and analysis steps of time course and UGP2 KO scRNA-seq data were run in Python 3 using Scanpy v.1.4.6+^20^ and anndata v.0.7.1+^49^ except stated otherwise. Downstream analysis of the scRNA-seq data of the mTOR inhibition experiments was performed in Seurat v5^50^. All subsequent preprocessing and analysis steps of scATAC-seq data were run in R v4.1.2+ using ArchR v1.0.2+^51^ except stated otherwise. All scRNA-seq and scATAC-seq figures were plotted using matplotlib, seaborn and ggplot2.

#### Species assignment and quality control

To mitigate technical cross-species batch effects, we pooled an equal number of cells from the three species for each of the samples. Consequently, we aligned resulting sequencing data against each of the three respective reference genomes separately. For time course single cell multiome sequencing and scRNA-seq of the UGP2 KO, we employed a two-step approach to assign each barcode to the species of origin: initially, we prematurely assigned cells to a species based on which reference genome yielded the highest counts per cell. Subsequently, we used souporcell^52^ to identify and remove doublet cells, and cluster cells on their genotype and their respective species of origin based on single nucleotide polymorphisms. For each sample, we filtered barcodes to retain high quality cells based on the total distributions of unique molecular identifier counts and genes. Barcodes with a fraction of mitochondria-encoded genes over 30% were excluded, likely indicating dying or stressed cells. Finally, we excluded genes detected in fewer than 20 cells from further analyses. For time course multiome data, UMI counts of each cell were normalized using the SCRAN algorithm as implemented in the R-based package^53,54^. Briefly, size factors that correlate with the amount of counts of captured cells were estimated and used for normalization before log-transforming the data. For UGP2 KO data, Scanpy’s function sc.pp.normalize_total was used to normalize the data. For scRNA-seq data of mTOR inhibition experiments, only cells with a minimum feature count of > 3000 and a maximum mitochondrial gene percentage of < 15% were considered.

For the time course scATAC-seq data, barcodes from the respective species were assigned using the labeled cell barcodes from the scRNA-seq analysis. Peaks were called using Macs2^55^. Next, we inspected the resulting peak and tile matrices based on the number of fragments generated from Tn5 enzyme transposition events, transcription start site (TSS) enrichment score and nucleosome signal per cell to obtain high quality cells. We excluded cells with <1,000 captured fragments or TSS enrichment scores <1 from further analysis. Additionally, we excluded peaks located on non-standard chromosomes or chromosome scaffolds, as well as peaks within genomic blacklist regions as defined by the ENCODE consortium^56^ from further analysis.

#### Feature selection and low dimensional embedding

We identified the top 4,000 variable genes based on normalized dispersion^57^ as adopted in Scanpy (pp.highly_variable_genes). Briefly, genes were ordered along their mean expression before selecting the genes with the highest variance-to-mean ratio. We then performed principal-component analysis dimension reduction by computing 15 principal components on highly variable genes using Scanpy’s pp.pca. Next, a neighborhood graph was computed on the first 50 principal components using Scanpy’s pp.neighbors with 15 neighbors. To identify genes involved in the linear NPC differentiation process, we used Waddington Optimal Transport (WOT)^58^ to identify potential driver genes correlating with fate probabilities towards the terminal macrostate as implemented in cellrank^59^. WOT, primarily used to infer developmental trajectories by minimizing transportation costs between cell states for timelapsed scRNA-seq data, was employed here to identify genes that follow the linear trajectory towards endpoint cells at the final time point. For two-dimensional visualization, we embedded the neighborhood graph via uniform manifold approximation and projection (UMAP)^60^ on these lineage driver genes identified from WOT with an effective minimum distance between embedded points of 0.5.

For the scATAC-seq data, we performed layered dimensionality reduction using latent semantic indexing (LSI), consisting of normalization via term frequency-inverse document frequency (TF-IDF) and dimension reduction via singular value decomposition (SVD). Finally, we calculated a UMAP embedding based on the LSI reduced dimensions with 30 neighbors and an effective minimum distance between embedded points of 0.5. To obtain reliable cell annotations, we transferred labels from scRNA-seq barcodes to scATAC-seq barcodes.

#### Simulated gene expression

To facilitate the visualization and interpretation of scATAC-seq data, we leveraged chromatin accessibility patterns to estimate gene expression profiles for cell state-specific marker genes. We calculated gene scores estimating the level of gene expression based on the local accessibility of the gene region, including the promoter and gene body, across all cells in the data, adjusting for gene distances and large differences in gene size using the GeneScoreMatrix in ArchR^51^. To further enhance the visualization, we used MAGIC^61^ to denoise gene activity scores.

#### Ingest mapping

To investigate the overall speed of differentiation between species and the effect of mTOR inhibition and loss of UGP2 in pluripotent stem cells across the three species, we used Scanpy’s ingest function (sc.tl.ingest). For species comparison of time course gene expression data from scRNA-seq and gene expression prediction data from scATAC-seq, we mapped cells from human, mouse and macaca into one common embedding. Integration of embeddings and annotations of macaca and human cells via a k-NN classifier was done, with mouse cells serving as the reference since they exhibited the fastest differentiation speed and therefore covered the most stages during the differentiation process. To investigate the effect of mTOR inhibition and loss of UGP2 we used the respective species’ time course data as the reference for mapping. For the mTOR inhibition data, the proportion of cells assigned to a specific time point of the reference data was quantified using a custom-made pheatmap-based function^47^.

#### Linear regression time mapping

To explore the correlation between the mapped time derived from ingest mapping for scRNA and scATAC-seq and the corresponding sampling time points across the human and cynomolgus datasets, linear regression analysis was employed using the scikit-learn tool v1.5.0^62^. Mean values obtained over the mapped time were calculated for each dataset to enable a comparative examination. The regression models were constrained to pass through the origin. The fit for each model was evaluated using the R-squared statistic. Relative differentiation speeds were determined by calculating the slopes of the linear fits relative to mouse.

#### Differential gene expression and enrichment analysis

For differential gene expression analysis, Scanpy’s sc.tl.rank_genes_groups was applied to compare gene expression levels between previously defined clusters using the Wilcoxon method, aiming to identify genes that are significantly upregulated or downregulated in one cluster compared to others. After identifying differentially expressed genes (DEGs), enrichment analysis was conducted. Lists of DEGs were analyzed using Gene Ontology (GO) or Kyoto Encyclopedia of Genes and Genomes (KEGG) databases, indicating potential biological processes or pathways that are active in one cell population over another.

#### Clustering and identification of fast- and slow-differentiating cells within each species

The Leiden algorithm was utilized for unsupervised clustering for all species. Subsequently, the tl.rank_genes_groups method was applied within Leiden clusters, employing the Wilcoxon test to identify DEGs. The resulting DEG lists were exported for each species and further refined by retaining only those genes that were common across all three species and displayed a p-value < 0.01 and a log-fold change > 2. To check for correct assignment, the clusters were divided into smaller subclusters and each subcluster was reevaluated and checked for correct assignment. Again, the tl.rank_genes_groups method was applied to determine DEGs between refined clusters and only genes common to all three species and displaying a p-value < 0.01 and a log-fold change > 2 were retained. These genes were used as marker genes for the clusters pluripotency high - pluripotency low - intermediate - neural low - neural high and neuronal within the dataset of each species. Using the sc.tl.score_genes function of Scanpy, we calculated gene scores for each cell and each marker gene set defining the respective cluster. Further subdivision was carried out, and smaller subclusters were analyzed for overlap with the marker genes. For the identification of fast- and slow-differentiating cells, we determined the mean gene score for each cluster and used this value to set a threshold to select single cells that most confidently reflected the respective gene expression profiles. These subgroups of cells were further subdivided according to the time point at which they reached their specific differentiation state. For a single differentiation state, we focused on the two time points that contained most of the cells in that differentiation state, and defined the cells found at the earlier of the two time points as fast-, and the cells found at the later time-point as slow-differentiating.

## Author contributions

A. S., C.M., M.D., A.P., J.S., and M.T. contributed to the conceptualization of the study. A.P. and J.S. conducted the laboratory experiments, and M.T. conducted the computational investigations. M.T., A.P., and J.S. performed data analysis. J.G. and W.E. generated and provided cyiPSCs, L.E.S. and T.S.B. generated and provided H9 UGP2 KO lines, M.S. conducted library preparations for time course single-cell multiome sequencing. X.P. conducted pre-processing and demultiplexing of the time course single-cell multiome sequencing data and UGP2 KO scRNA-seq data. A.P., J.S., and M.T. carried out visualization of the data. A.P., J.S. and M.T. wrote the original draft of the manuscript, with C.S., C.M., and M.D. contributing to the review and editing. Supervision was provided by C.S., C.M., and M.D. Funding acquisition was handled by C.S., C.M., and M.D.

## Acknowledgements

We want to thank the members of the Schröter, Marr and Drukker labs for the fruitful discussions and their valuable feedback over the years, with special thanks to Max Fernkorn for handling the GEO data upload. We thank Ludovic Vallier (MPI for Molecular Genetics) and Derk ten Berge (Erasmus MC) for providing the mEpiSCs, and the Studer laboratory for the PAX6 reporter. J.S. and C.S. thank Philippe Bastianes and the Department of Systemic Cell Biology at the MPI Dortmund for support and discussions. We extend our appreciation to Stefano Maffini and Department 1 for their assistance with nucleofection protocols, and to Shamya Noeen for her support with cell tracking. Our thanks also goes to Antonio Scialdone for the critical evaluation and internal review. We acknowledge the technical support of Core Facility Genomics at Helmholtz Munich. Lastly, we appreciate the technical expertise and assistance provided by Ejona Rusha and Anna Pertek from the iPSC Core Facility at Helmholtz Munich throughout this project. We acknowledge support from the Hightech Agenda Bayern. This work was funded by the VW foundation (project no. A130140 “OntoTime”), the European Research Council (Horizon 2020 Research and Innovation Programme, grant agreements no. 866411 and 101113551), and the Max Planck Society. T.S.B. acknowledges ongoing support by the Netherlands Organization for Scientific Research (ZonMw Vidi, grant 09150172110002), a Citizens United for Research in Epilepsy (CURE) Epilepsy Award 2021, and funds from EpilepsieNL (WAR 2022-01).

## Data and code availability

Raw data and count matrices of scRNA-seq data and scATAC-seq data are available on GEO (accession number: GSE275572). Further processed sequencing data, as well as custom analysis scripts will be made available upon reasonable request.

## Supplementary Figures

**Figure S1:**
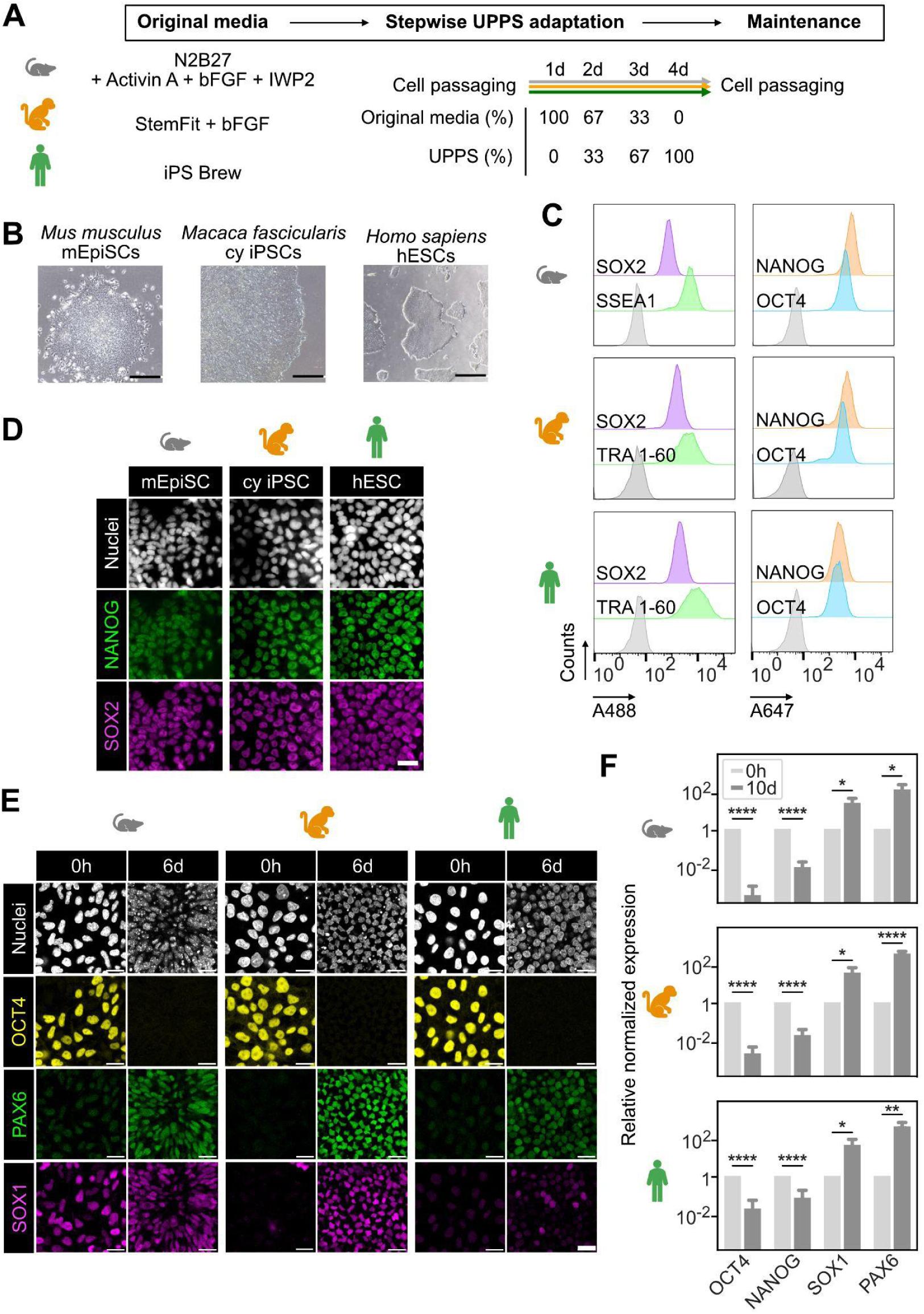
Primed epiblast-like pluripotent stem cells of three mammalian species cultured and differentiated under identical conditions. A. Schematic illustrating the stepwise adaptation process to harmonized UPPS medium from the original culture conditions of mEpiSC (top), cyiPSC (middle) and hESC (bottom). B. Brightfield images of mouse (left), cynomolgus (middle) and human (right) pluripotent stem cells cultured in harmonized conditions using Matrigel and UPPS. Cells from all three species display regular colony morphology typical for pluripotent cells. Scale bars, 500 µm. C. Flow cytometric analysis for pluripotency marker expression in mEpiSCs (top), cyiPSCs (middle) and hESCs (bottom) cultured in harmonized conditions using Matrigel and UPPS. Left panels show distribution of SOX2 (purple) and SSEA1/TRA 1-60 (green) expressions, right panels show NANOG (orange) and OCT4 (blue) expressions. Gray distributions represent unstained control samples. D. Immunofluorescence staining of pluripotency markers NANOG (green) and SOX2 (magenta) in mEpiSCs (top), cyiPSCs (middle) and hESCs (bottom) cultured in harmonized conditions using Matrigel and UPPS. Scale bars, 25 µm. E. Immunofluorescence staining of the pluripotency marker OCT4 (yellow) and the neural markers PAX6 (green) and SOX1 (magenta) in mEpiSCs (left), cyiPSCs (middle) and hESCs (right) before and after 6 days of NPC differentiation. Scale bars, 20 µm. F. RT-qPCR analysis of relative expression level of the pluripotency markers OCT4 and NANOG, and neural markers SOX1 and PAX6 in mEpiSCs (top), cyiPSCs (middle) and hESCs (bottom) after 10 days of NPC differentiation (dark bars). Expression levels were normalized for each gene to its expression in pluripotency conditions (light bars) using the 2^-ddCt method. Error bars indicate standard deviation (N = 3). Significance was was determined using the Student’s t-test: * p < 0.05, ** p < 0.01, *** p < 0.001, **** p < 0.0001.

**Figure S2:**
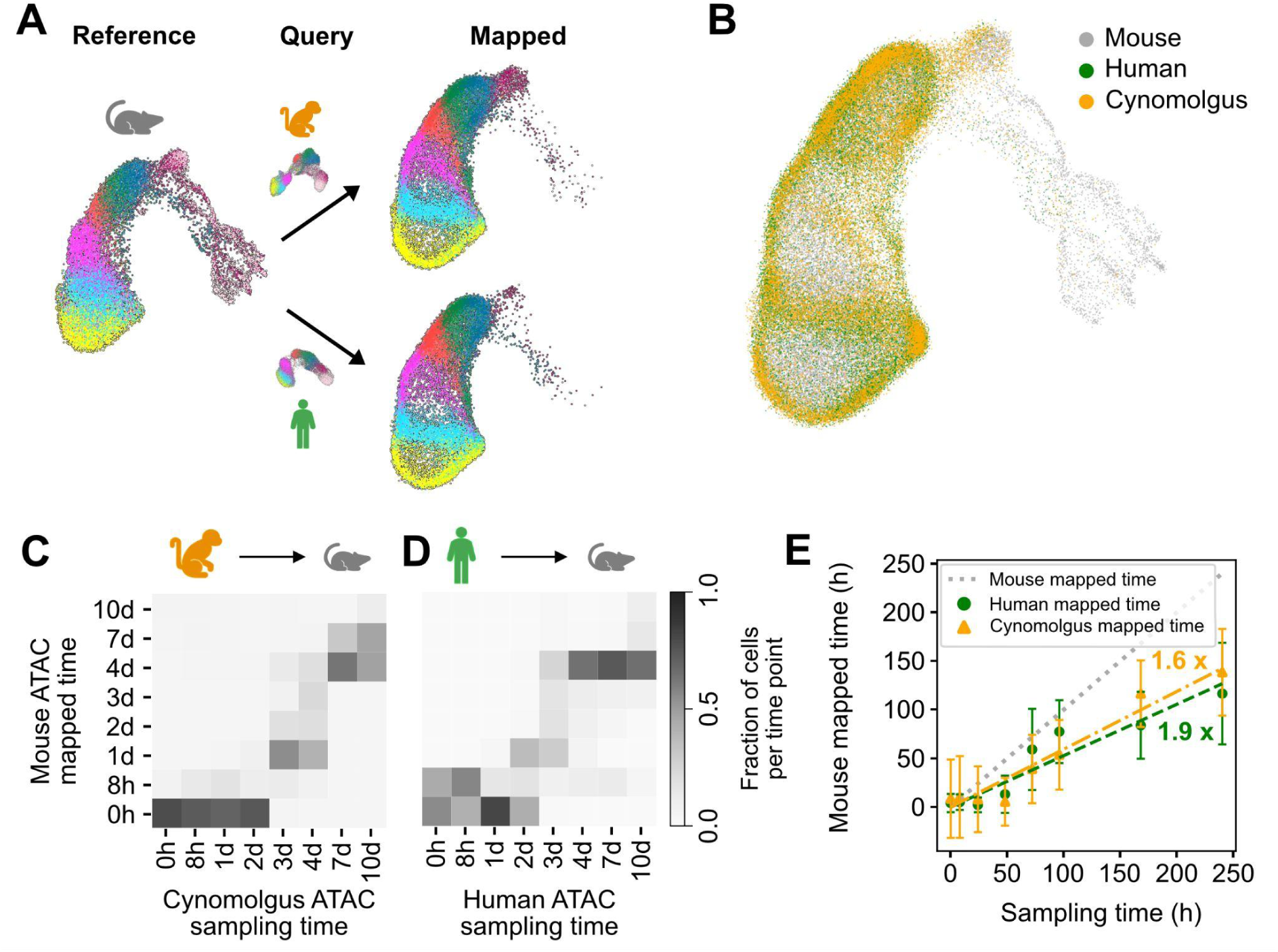
Inter-species single-cell mapping reveals that mouse cells differentiate fastest, followed by cynomolgus and human on RNA and chromatin level. A. Schematic illustrating approach to map single-cell transcriptomes from human and cynomolgus cells along the differentiation time-course onto the mouse reference embedding. B. Common embedding of human (green) and cynomolgus cells (yellow) in the mouse reference UMAP map (gray). Neither human nor cynomolgus cells reached the most differentiated states seen in the mouse, but the most advanced cynomolgus cells were slightly further along in the differentiation trajectory compared to human cells, indicating a marginally faster differentiation pace in cynomolgus compared to human. C-D. Heatmaps using simulated gene expression from scATAC-seq in cynomolgus and human cells compared to simulated gene expression from scATAC-seq in mouse cells. Tile colors indicate fractions of cynomolgus (C) and human (D) cells for each sampling time mapping to reference time points in mouse. E. Linear regression of cross-species mapping using simulated gene expression determined from scATAC-seq. Data points show the mean mapped time of human (green circles) and cynomolgus (orange triangles) cells on the mouse reference data. Lines are linear fits to the data for human (green, y = 0.53x, R^2^ = 0.89) and cynomolgus (orange, y = 0.59x, R^2^ = 0.95), and indicate 1.7 and 1.9-fold faster differentiation in mouse compared to cynomolgus and human, respectively, when determined using chromatin dynamics.

**Figure S3:**
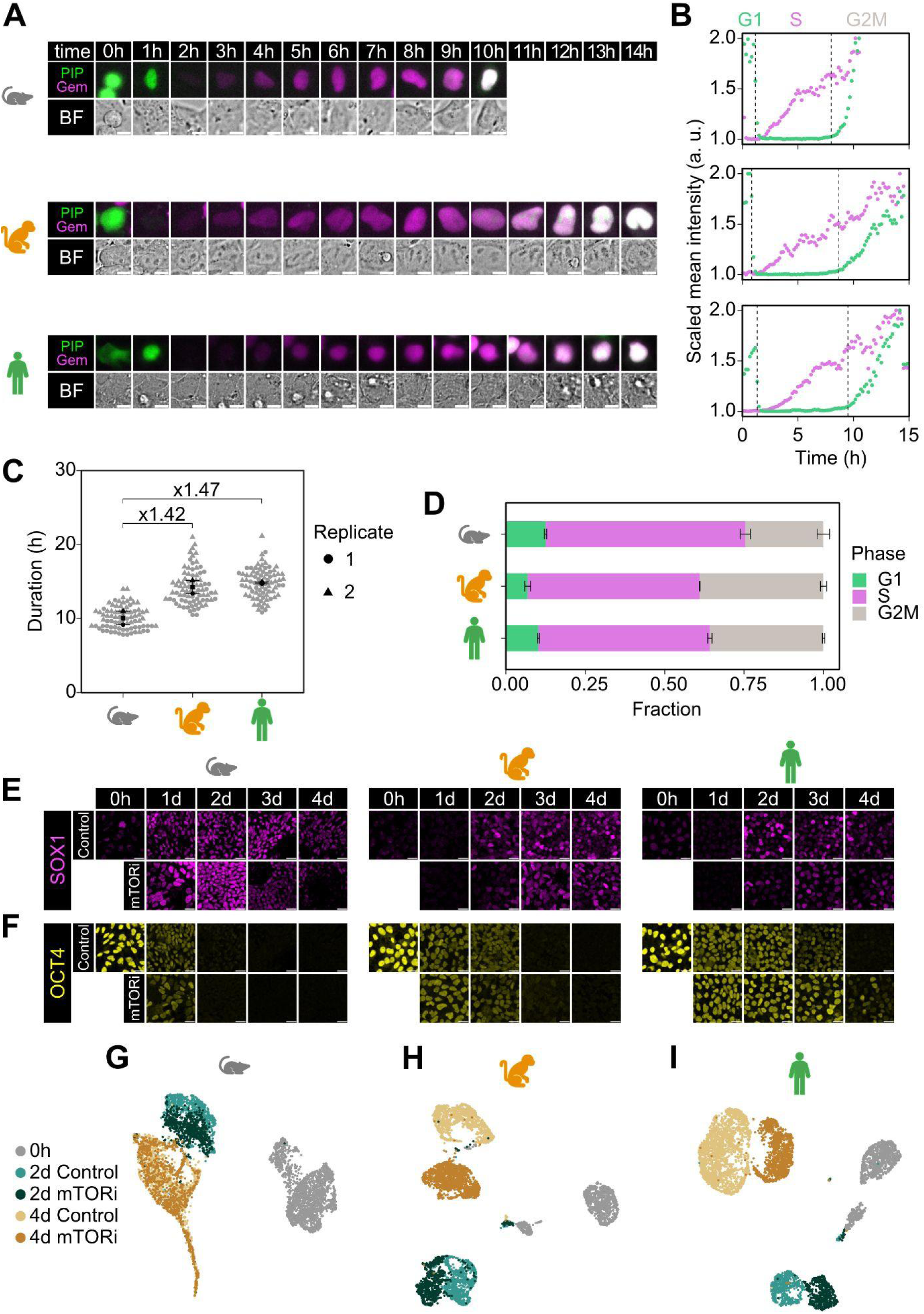
Cell cycle durations are species-specific but growth and cell cycle manipulation via mTOR inhibition does not delay early neural differentiation. A. Stills from movies of mEpiSCs (top), cyiPSCs (middle) and hESCs (bottom) expressing the PIP-FUCCI sensor. Expression of Cdt_1-17_-NLS-HA-mVenus (PIP) in green, expression of mCherry-Gem_1-110_ (Gem) in magenta. Scale bars, 10 µm. B. Fluorescence profiles derived from cells shown in A. Dashed lines indicate transitions between the cell cycle phases G1, S and G2M (See methods for criteria to determine cell cycle phase transitions). C. Distributions of cell cycle durations in mEpiSCs (left), cyiPSCs (middle) and hESCs (right). Data from two independent experiments indicated by different shapes. Light symbols show data from individual cells, dark circles and triangles show mean of each replicate, black squares indicate average of replicates, bars indicate standard error. Cell cycle durations are broadly distributed within each species, but average cell cycle durations are species-specific. D. Cell cycle phase durations normalized to total cell cycle length. Cells from all species show a similar cell cycle structure characteristic for pluripotent cells with a short G1-phase (green) and dominant S- and G2/M-phases (magenta and gray). E-F. Same differentiation time course as in Figure 3C showing mouse (left), cynomolgus (middle) and human cells (right) fixed after 0, 1, 2, 3, and 4 of NPC differentiation in the absence (control) or presence of mTOR inhibition, but stained for the neural marker SOX1 (magenta, E), and the pluripotency marker OCT4 (yellow, F). mTORi has no discernable effect on SOX1 expression onset, but leads to a slightly longer maintenance of OCT4 expression in human cells, possibly due to reduced cell proliferation and protein dilution. Scale bars, 20 µm. G-I. UMAP representation of scRNA-seq data from mEpiSCs (G), cyiPSCs (H) and hESCs (I) differentiated for 2 (green) or 4 days (orange) in the presence (dark symbols) or absence (control, light symbols) of the mTOR-inhibitor INK128. Transcriptomes of undifferentiated cells are shown in gray. mTOR-inhibited and control cells are overlaid in mouse but map away from each other in the primate species, consistent with stronger effects of the inhibitor in the primate compared to mouse cells.

**Figure S4:**
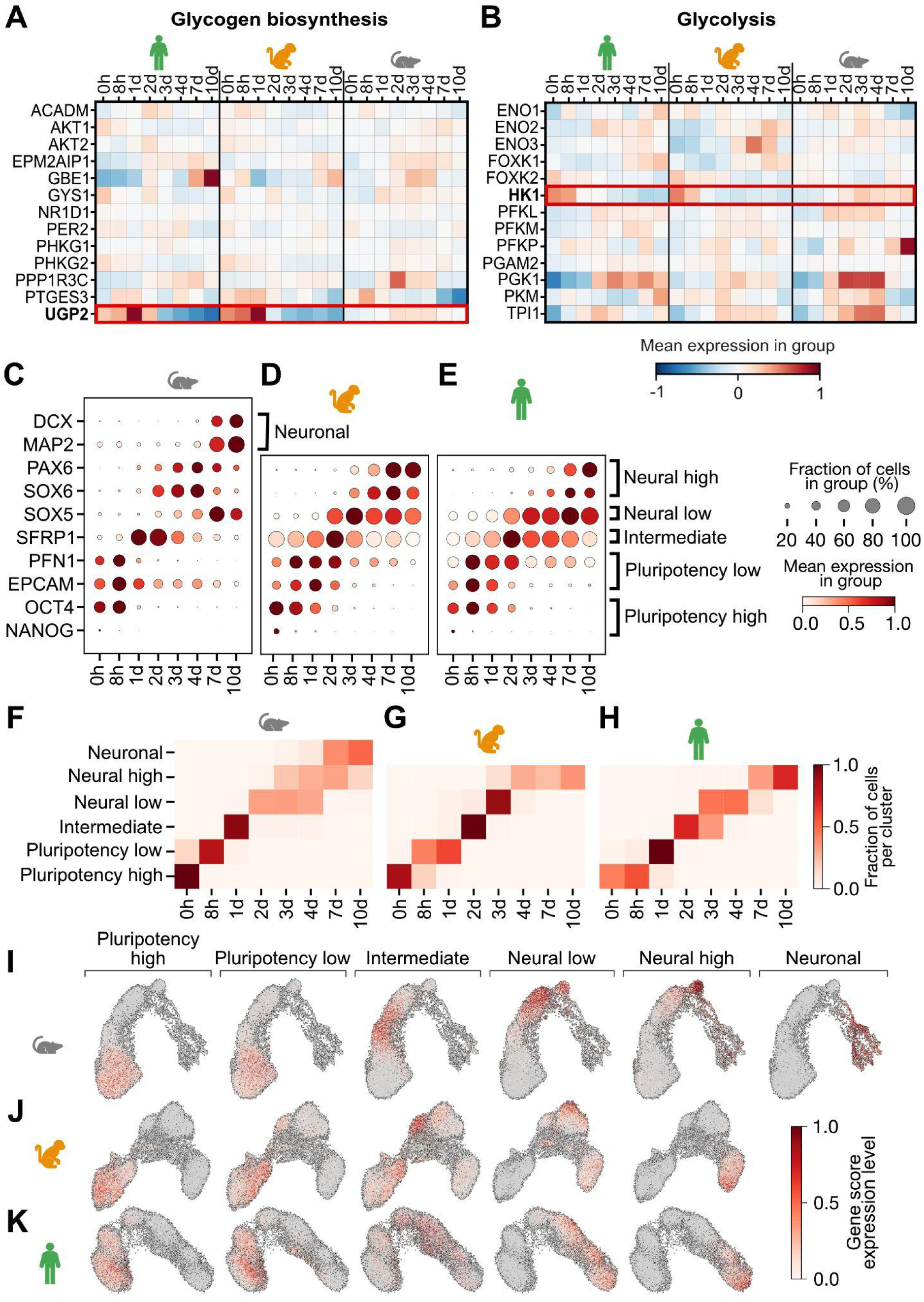
Identification of candidate genes for differentiation speed through comparison of fast- and slow-differentiating cells within and across species. A-B. Heatmap showing the mean normalized expression of individual genes associated with glycogen biosynthesis (A) and glycolysis (B) across the differentiation time-course in human (left), cynomolgus (middle), and mouse cells (right). Red box highlights UGP2 (A) and HK1 (B) expression which differ most strongly between human, cynomolgus, and mouse. C-E. Expression of selected marker genes used to annotate cells as pluripotency high, pluripotency low, intermediate, neural low, neural high and neuronal during NPC differentiation of mouse (C), cynomolgus (D) and human (E) cells. Marker genes are expressed in the same relative sequence across species, but at later absolute times in the slower differentiating primates. F-H. Heatmap showing the fraction of cells with specific state annotations coming from different sampling time-points during NPC differentiation for mouse (F), cynomolgus (G) and human (H). The same differentiation states are reached at progressively later time-points in cynomolgus and human cells compared to the mouse. I-K. Expression scores of gene groups defining the differentiation states pluripotency high, pluripotency low, intermediate, neural low, neural high and neuronal (from left to right) shown on the UMAP embeddings from Figure 1. See Methods for choice of gene groups and calculation of expression scores.

**Figure S5:**
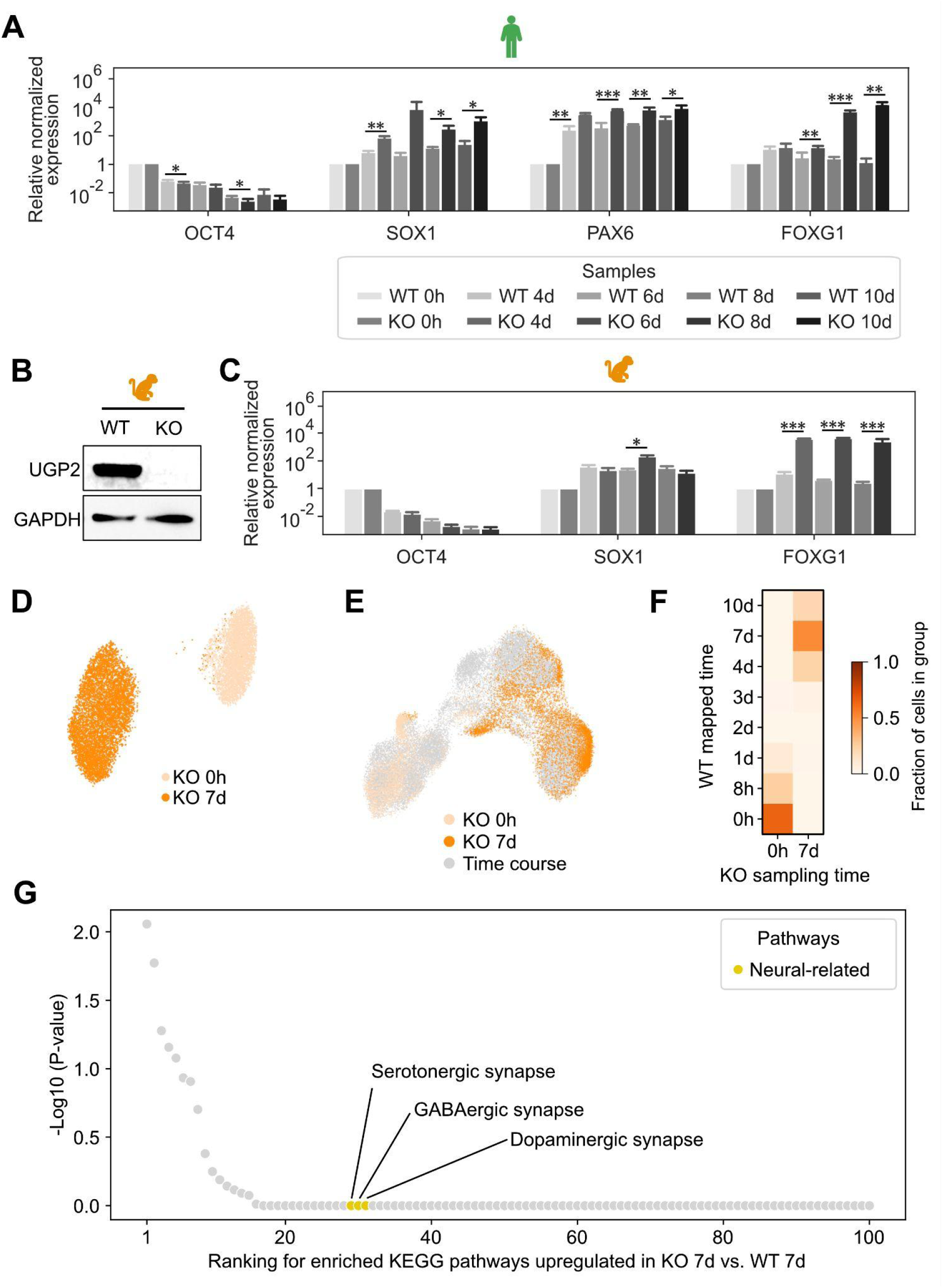
Impact of UGP2 knockout on neural differentiation in hESCs and cyiPSCs. A.-B. RT-qPCR expression analysis of the pluripotency marker OCT4, the early neural markers SOX1 and PAX6, and later-stage neural marker FOXG1 after 0, 4, 6, 8 and 10 days of NPC differentiation in wild-type (light) and UGP2 KO (dark) hESCs. Relative expression was calculated using the 2^-ddCt method and normalized using the 0h sample as reference. Data from N = 3 independent experiments, error bars represent standard deviation. Significance was determined using the Student’s t-test: * p < 0.05, ** p < 0.01, *** p < 0.001. Neural markers are consistently more highly expressed in UGP2 knockout compared to the wild-type cells. B. Western blot analysis of UGP2 protein in cynomolgus wild-type iPSCs (WT, reproduced from Figure 5C) and UGP2 knockout (KO) cells. UGP2 protein is absent in the cynomolgus iPSCs KO line. GAPDH is used as housekeeping control. C. RT-qPCR expression analysis as in (A) but comparing OCT4, SOX1 and FOXG1 in cynomolgus wild-type and UGP2 KO iPSCs until day 8 of differentiation. The later-stage marker FOXG1 is more highly expressed in KO compared to wild-type cells. D. UMAP representation of scRNA-seq data of cynomolgus UGP2 KO cells before (0h) and after 7 days of NPC differentiation. E. UMAP projection of single-cell transcriptomes of UGP2 KO cells at 0 hours (light orange) and 7 days (dark orange) of differentiation onto the time-resolved reference data set (gray). F. Heatmap showing the fraction of cynomolgus UGP2 KO cells at 0 hours and 7 days of NPC differentiation that map to specific time points of the cynomolgus wild-type reference data set. G. KEGG pathway enrichment using differentially expressed genes in cynomolgus UGP2 KO versus wild-type cells at day 7 of NPC differentiation. Top 100 picks sorted by p-value are displayed, neural related pathways are highlighted in yellow. In contrast to the situation in human cells, neural related pathways are not enriched in cynomolgus KO cells.

