## Supplementary material for "Single-cell multiome uncovers differences in glycogen metabolism underlying species-specific speed of development": Tables S2 - S4

**Supplementary Tables S2-S4**

**Table S2: Reagents and Resources**

| **REAGENT or RESOURCE** | **SOURCE** | **IDENTIFIER** |
| --- | --- | --- |
| **Antibodies** | | |
| Anti-Goat IgG (H+L) Cross-Adsorbed Secondary Antibody, Alexa Fluor™ 594 | Thermo Fisher Scientific | Cat#: A11058;  RRID: AB_142540 |
| Anti-Goat IgG (H+L) Cross-Adsorbed Secondary Antibody, Alexa Fluor™ 568 | Thermo Fisher Scientific | Cat#: A-11057;  RRID: AB_2534104 |
| Anti-Mouse IgG (H+L) Highly Cross-Adsorbed Secondary Antibody, Alexa Fluor™ 488 | Thermo Fisher Scientific | Cat#: A21202; RRID:AB_141607 |
| Anti-Mouse IgG (H+L) Highly Cross-Adsorbed Secondary Antibody, Alexa Fluor™ 647 | Thermo Fisher Scientific | Cat#: A-31571;  RRID: AB_162542 |
| Anti-Mouse IgG & IgM Antibody, HRP conjugate | Sigma Aldrich | Cat#: AP130P;  RRID: AB_91266 |
| Anti-Rabbit IgG (H+L) Highly Cross-Adsorbed Secondary Antibody, Alexa Fluor™ 647 | Thermo Fisher Scientific | Cat#: A-31573;  RRID: AB_2536183 |
| Anti-Rabbit IgG (H+L) Highly Cross-Adsorbed Secondary Antibody, Alexa Fluor™ 488 | Thermo Fisher Scientific | Cat#: A-21206;  RRID: AB_2535792 |
| Goat anti-SOX1 | R&D Systems | Cat#: AF3369;  RRID: AB_2239879 |
| Goat IgG isotype control | R&D Systems | Cat#: AB-108-C;  RRID: AB_354267 |
| Mouse anti-GAPDH | Thermo Fisher Scientific | Cat#: MA1-16757;  RRID: AB_568547 |
| Mouse anti-OCT4 | Santa Cruz | Cat#: sc-5279;  RRID: AB_628051 |
| Mouse anti-SOX2 | R&D Systems | Cat#: MAB2018; RRID:AB_358009 |
| Mouse anti-SSEA1 | Santa Cruz | Cat#: sc-21702;  RRID: AB_626918 |
| Mouse anti-TRA-1-60 | abcam | Cat#: ab16288;  RRID: AB_778563 |
| Mouse IgG isotype control | GeneTex | Cat#: GTX35009; RRID:AB_10618187 |
| Mouse IgM isotype control | Santa Cruz | Cat#: sc-3881;  RRID: AB_737292 |
| NANOG | Cell Signaling Technology | Cat#: 4903;  RRID: AB_10559205 |
| NANOG (mouse specific) | Cell Signaling Technology | Cat#: 8822;  RRID: AB_11217637 |
| Rabbit anti-OCT4 | GeneTex | Cat#: GTX101497;  RRID: AB_10618784 |
| Rabbit anti-PAX6 | BioLegend | Cat#: 901301; RRID:AB_2565003 |
| Rabbit anti-UGP2 | Santa Cruz | Cat#: sc-514174 |
| Rabbit IgG isotype control | GeneTex | Cat#: GTX35035; RRID:AB_10623175 |
| **Chemicals, peptides, and recombinant proteins** | | |
| β-Mercaptoethanol | Thermo Fisher Scientific | Cat#: 31350-010 |
| Accutase® solution | Sigma-Aldrich | Cat#: A6964 |
| Agarose | Biozym | Cat#: 840004 |
| B-27 Supplement (50x) | Thermo Fisher Scientific | Cat#: 17504044 |
| Blasticidin | Gibco | Cat#: R21001 |
| Bovine Albumin Fraction V, 7.5% solution | Thermo Fisher Scientific | Cat#: 15260037 |
| CHIR99021 | R&D Systems | Cat#: 4953/10 |
| Clarity Western ECL Substrate | BioRad | Cat#: 1705060S |
| Digitonin (5%) | Thermo Fisher Scientific | Cat#: BN2006 |
| Dimethyl sulfoxide, DMSO | Honeywell | Cat#: D5879 |
| DMEM/F12 | PAN Biotech | Cat#: P04-41250 |
| DNA Gel loading dye, 6x | Thermo Fisher Scientific | Cat#: R0611 |
| DPBS | Thermo Fisher Scientific | Cat#: 14190094 |
| DPBS with Ca^2+^/Mg^+^ | Thermo Fisher Scientific | Cat#: 14040091 |
| DTT | Thermo Fisher Scientific | Cat#: 18080044 |
| EDTA, 0.5 M | Promega | Cat#: V4231 |
| Formaldehyde 4%, ROTI Histofix | Carl Roth | Cat#: P087.1 |
| GlutaMAX (100x) | Thermo Fisher Scientific | Cat#: 35050-038 |
| Glycerol | Gerbu | Cat#: 2006.5000 |
| Human Recombinant Insulin (4 mg/mL) | Thermo Fisher Scientific | Cat#: 12585014 |
| INK128 | Cell Signaling | Cat#: 30690 |
| IWR-1 | Sigma-Aldrich | Cat#: I0161 |
| Laemmli Buffer, 2x | BioRad | Cat#: 1610737EDU |
| LDN-193189 HCl | Peprotech | Cat#: 1066208 |
| Lipofectamine2000 | Thermo Fisher Scientific | Cat#: 11668019 |
| Magnesium Chloride solution, 1 M | Sigma-Aldrich | Cat#: M1028 |
| Matrigel® Matrix | Corning | Cat#: 354234 |
| MEM Non-Essential Amino Acids Solution (100x) | Thermo Fisher Scientific | Cat#: 11140-035 |
| Methanol | Serva Electrophoresis | Cat#: 45631.02 |
| Mini PROTEAN® TGX stain-free gel | BioRad | Cat#: 4568023 |
| n-propyl-gallate | Sigma Aldrich | Cat#: 02370-100G |
| N2 Supplement (100x) | Thermo Fisher Scientific | Cat#: 17502048 |
| NaCl (Sodium chloride) | Carl Roth | Cat#: P029.2 |
| Neuropan | PAN Biotech | Cat#: P04-00900 |
| NotI-HF | New England Biolabs | Cat#: R3189 |
| Powdered milk, blotting grade | Sigma Aldrich | Cat#: T145.1 |
| Protector RNase inhibitor | Sigma-Aldrich | Cat#: 3335402001 |
| Puromycin | Sigma Aldrich | Cat#: P8833 |
| RIPA Lysis and Extraction Buffer | Thermo Fisher Scientific | Cat#: 89900 |
| RNase-Free DNase Set | Qiagen | Cat#: 5000650 |
| ROCK inhibitor Y-27632 | R&D Systems | Cat#: 1254/10 |
| SB431542 | Peprotech | Cat#: 3014193 |
| Sodium Chloride solution, 5 M | Sigma-Aldrich | Cat#: 59222C |
| Sodium Dodecyl Sulfate | Serva Electrophoresis | Cat#: 20768.02 |
| StemMACS iPS Brew XF | Miltenyi Biotech | Cat#: 130-104-368 |
| SYBR Safe DNA GEL Stain | Thermo Fisher Scientific | Cat#: S33102 |
| TERGITOL™-Solution, Type NP-40 | Sigma-Aldrich | Cat#: NP40S |
| TRIS base | Carl Roth | Cat#: 5429.3 |
| Tris/Glycine Buffer (10x) | BioRad | Cat#: 1610734 |
| Triton™ X-100 | Sigma-Aldrich | Cat#: X100-500ML |
| Trizma Hydrochloride solution, pH 7.4 | Sigma-Aldrich | Cat#: T2194 |
| TrueCut™ Cas9 Protein v2 | Thermo Fisher Scientific | Cat#: A36496 |
| Trypan Blue Stain (0.4%) | Thermo Fisher Scientific | Cat#: T10282 |
| Tween-20 | Sigma-Aldrich | Cat#: P9416 |
| XhoI | New England Biolabs | Cat#: R0146 |
| **Critical commercial assays** | | |
| 3’ Feature Barcode Kit | 10x Genomics | Cat#: 1000262 |
| Chromium Next GEM Chip J Single-cell Kit | 10x Genomics | Cat#: 1000230 |
| Chromium Next GEM Single Cells 3’ Reagent Kits v3.1 (Dual Index) | 10x Genomics | Cat#: 1000128 |
| Chromium Next GEM Single-cell Multiome ATAC + Gene Expression Reagent Bundle | 10x Genomics | Cat#: 1000285 |
| DreamTaq Green DNA Polymerase (5 U/μL) | Thermo Fisher Scientific | Cat#: EP0712 |
| Glycogen Assay Kit | Abcam | Cat#: ab65620 |
| Inside Stain Kit | Miltenyi Biotech | Cat#: 130-090-477 |
| NovaSeq 6000 S1 Reagent Kit (100 cycles) | Illumina | Cat#: 20028319 |
| NovaSeq 6000 SP Reagent Kit (100 cycles) | Illumina | Cat#: 20028401 |
| P3 Primary Cell 4D-Nucleofector® X Kit | Lonza | Cat#: V4XP-3024 |
| Pierce™ BCA Protein Assay Kit | Thermo Fisher | Cat#: 23225 |
| QIAamp DNA Mini Kit | Qiagen | Cat#: 51304 |
| QIAquick PCR Purification Kit | Qiagen | Cat#: 28104 |
| RNeasy Mini Kit | Qiagen | Cat#: 5001329 |
| SYBR Green qPCR Master Mix | Thermo Fisher Scientific | Cat#: 4367659 |
| Verso cDNA Synthesis Kit | Thermo Fisher Scientific | Cat#: AB1453A |
| **Deposited data** | | |
| Cynomolgus genome GCA_011100615.1 (Genome assembly 6.0) | ENSEMBL | https://www.ensembl.org/Macaca_fascicularis/Info/Index |
| Human reference genome GRCh38 | Genome Reference Consortium | https://www.ncbi.nlm.nih.gov/datasets/genome/GCF_000001405.26/ |
| Mouse genome GRCm38 mm10 | Genome Reference Consortium | https://www.ncbi.nlm.nih.gov/datasets/genome/GCF_000001635.26/ |
| Mouse reference genome GRCm39 mm39 | Genome Reference Consortium | https://www.ncbi.nlm.nih.gov/datasets/genome/GCF_000001635.27/ |
| Sequencing data | Gene Expression Omnibus (GEO) | GSE275572 |
| **Experimental models: Cell lines** | | |
| *M. fascicularis*/Cynomolgus: cyiPSC | Radmer et al.^44^ | https://pubmed.ncbi.nlm.nih.gov/37590551/ |
| *M. musculus*/Mouse: EpiSC | Kurek et al.^42^ | https://pubmed.ncbi.nlm.nih.gov/25544567/ |
| *H. sapiens*/Human: H9 ESC | WiCELL Research Institute | https://www.wicell.org |
| *H. sapiens*/Human: H9 ESC PAX6::H2B-GFP | Tchieu et al.^25^ | https://www.ncbi.nlm.nih.gov/pmc/articles/PMC5737635/ |
| **Oligonucleotides** | | |
| Primers | Sigma-Aldrich | Table 1 |
| Guide RNAs | Thermo Fisher Scientific | Table 1 |
| **Recombinant DNA** | | |
| pBASE plasmid | Wang et al.^45^ | https://pubmed.ncbi.nlm.nih.gov/18579772/ |
| PiggyBAC H2B-Cerulean-IRES-bsr | Schumacher et al.^47^ | https://pubmed.ncbi.nlm.nih.gov/38871742/ |
| PiggyBAC vector | Wang et al.^45^ | https://pubmed.ncbi.nlm.nih.gov/18579772/ |
| pENTR-PIP-FUCCI | Grant et al.^21^ | https://pubmed.ncbi.nlm.nih.gov/30421640/  Addgene plasmid #118621 |
| **Software and algorithms** | | |
| Anndata v.0.7.1+ | Virshup et al.^49^ | https://github.com/scverse/anndata |
| ArchR v1.0.2+ | Granja et al.^51^ | https://github.com/GreenleafLab/ArchR |
| CellRanger ARC v2.0.0 | 10x Genomics | https://www.10xgenomics.com/support/software/cell-ranger-arc/latest |
| CellRanger v7.2.0 | 10x Genomics | https://www.10xgenomics.com/support/software/cell-ranger/latest |
| FlowJo | BD Biosciences | https://flowjo.com |
| ggplot2 v3.3.6 |  | https://github.com/tidyverse/ggplot2 |
| ImageJ v2.9.0 | NIH | https://imagej.net/software/fiji/downloads |
| Macs2 | Zhang et al.^55^ | https://github.com/macs3-project/MACS |
| Matplotlib |  | https://matplotlib.org |
| MicroManager 2.0 | Edelstein et al.^48^ | https://micro-manager.org/Version_2.0 |
| Python 3 | Python Software Foundation | https://www.python.org/ |
| Quantstudio | Thermo Fisher Scientific | https://www.thermofisher.com/de/en/home/technical-resources/software-downloads/quantstudio-12k-flex-real-time-pcr-system.html |
| R v4.1.2+ | The R Foundation | https://www.r-project.org/ |
| Scanpy v.1.4.6+ | Wolf et al.^20^ | https://github.com/scverse/scanpy |
| Scikit-learn v1.5.0 | Pedregosa et al.^62^ | https://scikit-learn.org/stable/ |
| SCRAN | L. Lun et al.^53^, Lun et al.^54^ | https://github.com/elswob/SCRAN |
| Seaborn |  | https://seaborn.pydata.org |
| Seurat v5 | Hao et al.^50^ | https://satijalab.org/seurat/ |
| SnapGene viewer software v7.1.1 | Dotmatics | https://www.snapgene.com/snapgene-viewer |
| Souporcell v2.0+ | Heaton et al.^52^ | https://github.com/wheaton5/souporcell |
| StarDist 2D | Schmidt et al.^32^ | https://github.com/stardist/stardist |
| TrackMate v7.9.2 | Tinevez et al.^46^ | https://github.com/trackmate-sc/TrackMate |
| ZEN Microscopy Software | Zeiss | https://www.zeiss.com/microscopy/en/products/software/zeiss-zen-lite.html |

**Table S3: List of oligonucleotides**

| **Primers for RT-qPCR** | | | | | |
| --- | --- | --- | --- | --- | --- |
| **Target Gene** | | **Target Species** | | **Sequence** | |
| *ASCL1* | | Human | | F – CCAAGCAAGTCAAGCGACAG  R – TTGTGCGATCACCCTGCTTC | |
| *ASCL1* | | Mouse | | F – TGGACTTTGGAAGCAGGATG  R – TGCATCTTAGTGAAGGTGCCC | |
| *ASCL1* | | Cynomolgus | | F – AGGACTTTGAAAGCAGGGTGA  R – GACCCGAGCAAGAGCTTTCA | |
| *FOXG1* | | Human | | F – GGCAAGGGCAACTACTGGAT  R – CTGAGTCAACACGGAGCTGT | |
| *FOXG1* | | Mouse | | F – CTGATTGGTTCGGCAGTAGGA  R – TAGCAAAAGCTGCAACCACC | |
| *FOXG1* | | Cynomolgus | | F – GGCAAGGGCAACTACTGGAT  R – CTGAGTCAACACGGAGCTGT | |
| *GAPDH* | | Primates, mouse | | F – CATCACTGCCACCCAGAAGACTG  R – ATGCCAGTGAGCTTCCCGTTCAG | |
| *NANOG* | | Primates | | F – GATTTGTGGGCCTGAAGAAA  R – CAGATCCATGGAGGAAGGAA | |
| *NANOG* | | Mouse | | F – GAAATCCCTTCCCTCGCCAT  R – CAGGCATTGATGAGGCGTTC | |
| *OCT4* | | Primates | | F – GACAGGGGGAGGGGAGGAGCTAGG  R – CTTCCCTCCAACCAGTTGCCCCAAAC | |
| *OCT4* | | Mouse | | F – CCTGGGCGTTCTCTTTGGAA  R – ACCATACTCGAACCACATCCTTC | |
| *PAX6* | | Human | | F – CTAGCCAGGTTGCGAAGAAC  R – CTTGGGAAATCCGAGACAGA | |
| *PAX6* | | Mouse | | F – CTGAGGAACCAGAGAAGACAGG  R – CATGGAACCTGATGTGAAGGAGG | |
| *PAX6* | | Cynomolgus | | F – CCAAACAGAACTCTTGACAGGAA  R – TTCACTCCGCTGTGACTGTTC | |
| *SOX1* | | Human | | F – CATCTAGCGCCTTCGGGAC  R – AGTGCTTGGACCTGCCTTAC | |
| *SOX1* | | Mouse | | F – CGGATCTCTGGTCAAGTCGG  R – GGGACCTCGGTACAAAGTCG | |
| *SOX1* | | Cynomolgus | | F – CTGACGCATATCTAGCGCCT  R – GTGCTTGGACCTGCCTTACT | |
| *SOX2* | | Human | | F – AACCAGCGCATGGACAGTTA  R – GACTTGACCACCGAACCCAT | |
| *SOX2* | | Mouse | | F – CAAAAACCGTGATGCCGACT  R – CGCCCTCAGGTTTTCTCTGT | |
| *SOX2* | | Cynomolgus | | F – TTTGTCGGAGACGGAGAAGC  R – TAACTGTCCATGCGCTGGTT | |
| **Guide RNAs for CRISPR** | | | | | |
| **Gene** | **Target Species** | | **Label** | **Thermo ID** | **Sequence** |
| *UGP2* | Cynomolgus | | sgRNA_2 | GRWCZCZ | AAACUCAUGUGACGAUGCUG |
| *UGP2* | Cynomolgus | | sgRNA_3 | GRXGUWX | CGUCACAUGAGUUUGAGGUA |
| *UGP2* | Cynomolgus | | sgRNA_9 | GRYMNGV | AGCAAAGCAAUGUCUCAAGA |

**Table S4: List of antibodies and dilutions**

| **Target** | **Host** | **Dilution** | **Manufacturer** | **Cat. No.** |
| --- | --- | --- | --- | --- |
| **Primary Antibodies** |  |  |  |  |
| GAPDH | Mouse | 1:1000 | Thermo Fisher Scientific | MA1-16757 |
| NANOG | Rabbit | 1:100 | Cell Signaling Technology | 4903 |
| NANOG (mouse specific) | Rabbit | 1:100 | Cell Signaling Technology | 8822 |
| OCT4 | Rabbit | 1:100 | GeneTex | GTX101497 |
| OCT4 | Mouse | 1:100 | Santa Cruz | sc-5279 |
| PAX6 | Rabbit | 1:200 | BioLegend | 901301 |
| SOX1 | Goat | 1:200 | R&D Systems | AF3369 |
| SOX2 | Mouse | 1:100 | R&D Systems | MAB2018 |
| SSEA1 | Mouse | 1:20 | Santa Cruz | sc-21702 |
| TRA-1-60 | Mouse | 1:100 | abcam | ab16288 |
| UGP2 | Mouse | 1:1000 | Santa Cruz | sc-514174 |
| **Secondary Antibodies** |  |  |  |  |
| Anti-Goat IgG (H+L) Cross-Adsorbed Secondary Antibody, Alexa Fluor™ 594 | Donkey | 1:800 | Thermo Fisher Scientific | A-11058 |
| Anti-Mouse IgG (H+L) Highly Cross-Adsorbed Secondary Antibody, Alexa Fluor™ 488 | Donkey | 1:800 | Thermo Fisher Scientific | A-21202 |
| Anti-Mouse IgG & IgM Antibody, HRP conjugate | Goat | 1:10.000 | Sigma Aldrich | AP130P |
| Anti-Rabbit IgG (H+L) Highly Cross-Adsorbed Secondary Antibody, Alexa Fluor™ 647 | Donkey | 1:800 | Thermo Fisher Scientific | A-31573 |
| Anti-Rabbit IgG (H+L) Highly Cross-Adsorbed Secondary Antibody, Alexa Fluor™ 488 | Donkey | 1:500 | Thermo Fisher Scientific | A-21206 |
| Anti-Goat IgG (H+L) Cross-Adsorbed Secondary Antibody, Alexa Fluor™ 568 | Donkey | 1:500 | Thermo Fisher Scientific | A-11057 |
| Anti-Mouse IgG (H+L) Highly Cross-Adsorbed Secondary Antibody, Alexa Fluor™ 647 | Donkey | 1:500 | Thermo Fisher Scientific | A-31571 |
| **Isotype controls** |  |  |  |  |
| Goat IgG | Goat | variable | R&D Systems | AB-108-C |
| Mouse IgG | Mouse | variable | Thermo Fisher Scientific | 14471485 |
| Mouse IgM | Mouse | variable | Santa Cruz | sc-3881 |
| Rabbit IgG | Rabbit | variable | GeneTex | GTX35035 |
